# The lipid transfer function of RDGB at ER-PM contact sites is regulated by multiple interdomain interactions

**DOI:** 10.1101/2019.12.12.873810

**Authors:** Bishal Basak, Harini Krishnan, Padinjat Raghu

## Abstract

In *Drosophila* photoreceptors, following Phospholipase C-β activation, the phosphatidylinositol transfer protein (PITP) RDGB, is required to maintain lipid homeostasis at endoplasmic reticulum (ER) plasma membrane (PM) membrane contact sites (MCS). Depletion or mis-localization of RDGB results in multiple defects in photoreceptors. Previously, interaction between the FFAT motif of RDGB with the integral ER protein dVAP-A was shown to be important for its localization at ER-PM MCS. Here, we report that in addition to FFAT motif, a large unstructured region (USR1) of RDGB is required to support the RDGB/dVAP-A interaction. However, interaction with dVAP-A alone is insufficient for accurate localization of RDGB: this also requires association of RDGB with apical PM, through its C-terminal LNS2 domain. Deletion of LNS2 domain results in complete mis-localisation of RDGB and also induces large mis-regulated interdomain movements abrogating RDGB function. Thus, multiple independent interactions between individual domains of RDGB supports its function at ER-PM MCS.

## Introduction

The close approximation of intracellular membranes without fusion between them is emerging as a theme in cell biology (Gatta and Levine, 2017). Such apposition of membranes, referred to as membrane contact sites (MCS) can occur between multiple cellular organelles; most frequently, the endoplasmic reticulum (ER) which is the largest organelle, makes MCS with other cellular organelles including the plasma membrane (PM) (Cohen et al., 2018). ER-PM contact sites have been described in multiple eukaryotic cells, and are proposed to regulate a range of molecular process including calcium influx and the exchange of lipids (Chen et al., 2019; Saheki and Camilli, 2017).

The transfer of lipids between organelle membranes is a key function proposed for MCS. In the case of ER-PM contact sites, multiple lipids are thought to be transferred including phosphatidylserine (PS), phosphatidylinositol (PI), phosphatidic acid (PA), cholesterol and phosphatidylinositol 4-phosphate (PI4P) (Cockcroft and Raghu, 2018). These transfer activities are performed by several classes of lipid transfer proteins (LTPs). In order to carry out this function effectively, it is essential that these LTPs are accurately localized to ER-PM MCS, and several mechanisms that underlie this localization have been proposed (Alli-Balogun and Levine, 2019). LTPs frequently have multiple domains in addition to a lipid transfer domain. While some of these domains have been proposed to contribute to localization at the MCS, the *in vivo* function of several others is not clear. One group of LTPs named Phosphatidylinositol transfer proteins (PITPs) mediate specific transfer of PI and PA/Phosphatidylcholine (PC) between compartments. The first PITP identified and cloned by Dickenson and colleagues was a protein with a single phosphatidylinositol transfer domain (PITPd) (Dickeson et al., 1989). Since then multiple PITPs, with either single or multiple domains have been identified in various species (reviewed in Cockcroft *et al, 2010*). Importantly, in multi-domain PITPs, although the basic function of lipid transfer is conserved and restricted to the PITPd, the contribution of the additional domains to the regulation of PITPd activity *in vivo* is poorly understood.

*Drosophila* photoreceptors have emerged as an influential model system for the analysis of ER-PM contact sites (Yadav et al., 2016). The photoreceptors are polarized cells whose apical PM, also called rhabdomere, form contact sites with the sub-microvillar cisternae (SMC), a specialized domain of the photoreceptor ER [Fig 1A]. The apical PM and the SMC are specialized to mediate sensory transduction through G-protein coupled Phospholipase C-β (PLC-β) activation (Raghu et al., 2012). PLC-β activation triggers a series of enzymes whose substrates and products are lipid intermediates of the “PIP_2_cycle” (Cockcroft et al., 2016) that are distributed between the apical PM and the SMC. Some of these lipid intermediates such as PI and PA need to be transported between the apical PM and the SMC. *Drosophila* photoreceptors express a large multi-domain protein, **R**etinal **D**e**g**eneration **B** (RDGB) which has a well-annotated PITPd. Loss of function/hypomorphic mutants for *rdgB*, represented by *rdgB*^*2*^ and *rdgB*^*9*^ allelic flies respectively, show defective electrical responses to light, retinal degeneration and defects in light activated PIP_2_ turnover. The PITPd of RDGB has been shown to bind and transfer PI and PA *in vitro*, and is hence also sufficient to support RDGB function *in vivo* (Milligan et al., 1997; Yadav et al., 2015). Interestingly, the RDGB protein is localized exclusively to the MCS between the apical PM and the SMC (Vihtelic et al., 1993) [Fig 1A, B], thus offering an excellent *in vivo* setting to understand the relationship between LTP activity at an ER-PM contact site and it’s physiological function. RDGB is a large multidomain protein; the N-terminal PITPd is followed by several other domains including a FFAT motif, a large unstructured region (USR1), DDHD domain and LNS2 domain [Fig 1C-RDGB]. While the interaction of the FFAT motif with the ER integral protein, dVAP-A has been shown to be important for the localization and function of RDGB (Yadav et al., 2018), the relevance of the other domains and unstructured regions of RDGB to the function of the protein remains unknown.

**Figure 1:**
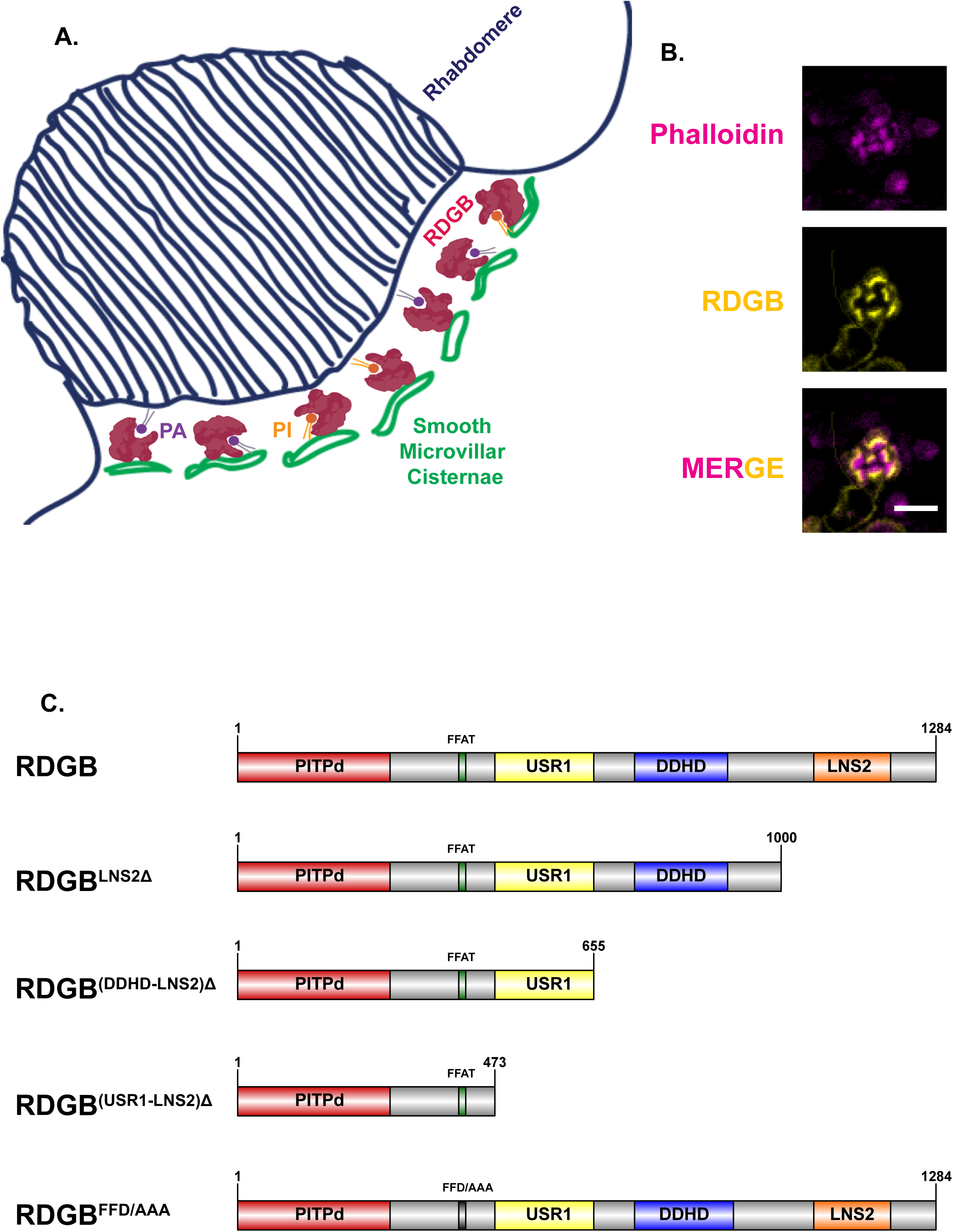
RDGB is a multidomain PITP acting at ER-PM contact site: **A.** Schematic of RDGB acting as a PITP at ER-PM contact site of *Drosophila* photoreceptor. The apical PM is thrown to numerous microvillar projections collectively termed as the rhabdomere, while the modified smooth ER compartment called Sub Microvillar Cisternae (SMC) is present at a distance of approximately ~10 nm from it. **B.** Confocal images of retinae obtained from wild type flies. Magenta represents phalloidin which marks the rhabdomeres and yellow represents RDGB. Scale bar= 5 µm. **C.** Domain structure of RDGB and the lists of constructs used in this study. RDGB protein is 1284 amino acid long and contains three domains-PITPd, DDHD and LNS2, a FFAT motif and a large unstructured region (USR1). Domains from C-terminal end of proteins were deleted to decipher functionally important regions of the protein. PITPd is marked in red, FFAT motif in green, USR1 in yellow, DDHD domain in blue and LNS2 domain in orange. The length of the protein is marked on each of the construct. RDGB^FFD/AAA^ represents the full length RDGB where each of the FFD residues in the FFAT motif has been mutated to alanine [Domain structure of RDGB drawn using Illustrator for Biological Sequences (IBS) software; http://ibs.biocuckoo.org/].

In this study, we report a role for two additional regions of the RDGB protein in supporting its normal localization and function. These regions, namely, the USR1 and the LNS2 domain are essential for interaction with the ER integral protein dVAP-A and the apical PM respectively, to enable correct localization of RDGB at the ER-PM MCS. Loss of the C-terminal domains, especially the LNS2 domain is associated with complete loss of protein function. Further, we find that these additional domains of RDGB interact with each other to mediate lipid homeostasis between the PM and the ER. These interactions, if perturbed, lead to functional consequences *in vivo*. Thus, this study demonstrates novel and key interdomain interactions for RDGB in mediating its lipid transfer activity at ER-PM MCS *in vivo*.

## Results

### The FFAT/dVAP-A interaction is not sufficient to stabilize the RDGB-dVAP-A protein complex

We tested the sufficiency of the FFAT/dVAP-A interaction for the localization of RDGB. For this, we reconstituted *rdgB*^*9*^ photoreceptors with a smaller RDGB protein containing the PITPd and the FFAT motif by truncating it just after the FFAT motif [Fig1C-RDGB^(USR1-LNS2)Δ^]. This truncated protein, RDGB^(USR1-LNS2)Δ^ was expressed in *rdgB*^*9*^ photoreceptors (*rdgB*^*9*^; GMR>*rdgB*^(USR1-LNS2)Δ^) [Fig 2A]. Since this protein has an intact PITPd and a FFAT motif, we hypothesized that RDGB^(USR1-LNS2)Δ^ would localize at the base of the rhabdomere similar to RDGB. However, surprisingly and in contrast to RDGB, we found that RDGB^(USR1-LNS2)Δ^ was mislocalized away from the base of the rhabdomere [Fig 2B]. To address why RDGB^(USR1-LNS2)Δ^ was mislocalized despite the presence of an intact FFAT motif, we tested if it could still interact with dVAP-A. For this, we immunoprecipitated endogenous dVAP-A using a dVAP-A specific antibody, and checked if RDGB co-immunoprecipitated along with it. We could pull down dVAP-A, and also detect wild type RDGB in the dVAP-A pulled down fraction, implying a physical interaction between the two proteins in wild type photoreceptors [Fig 2C]. This interaction was dependent on an intact FFAT motif; when the FFAT residues were mutated in full length RDGB [Fig 1C-RDGB^FFD/AAA^], it could not be immunoprecipitated with dVAP-A **[Fig 2D]**. Importantly, we found that RDGB^(USR1-LNS2)Δ^ could also not be immunoprecipitated with dVAP-A [Fig 2E]. Thus, while an intact FFAT motif is necessary for the interaction of RDGB with dVAP-A, it is not sufficient to support this interaction.

**Figure 2:**
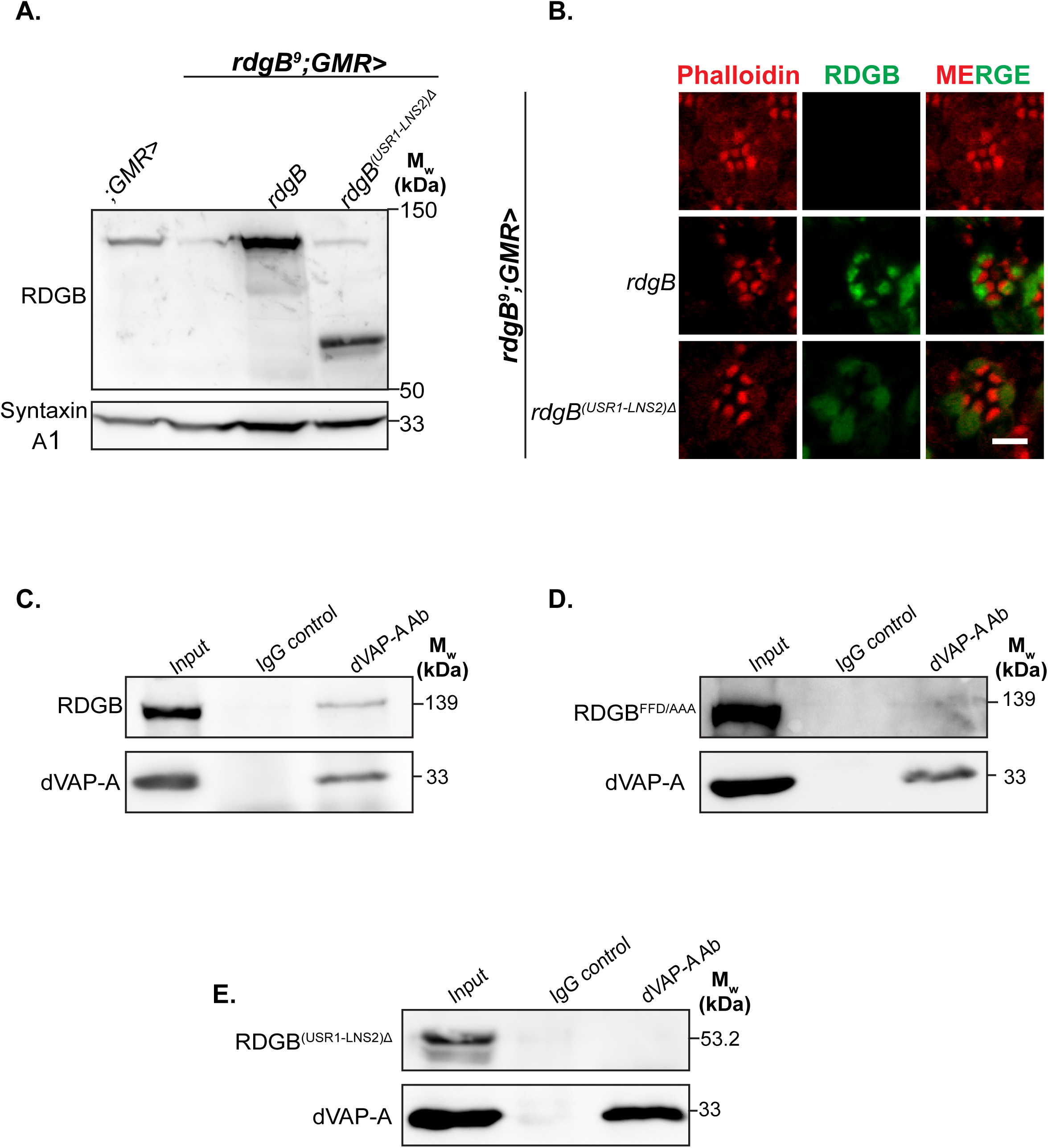
Interaction of RDGB with dVAP-A requires factors in addition to an intact FFAT motif. **A.** Western blot of head extracts made from flies of the mentioned genotype. The blot is probed with antibody to RDGB. SyntaxinA1 is used as a loading control (N=3). **B.** Confocal images of retinae obtained from flies of the mentioned genotypes. Red represents phalloidin which marks the rhabdomeres, and green represents RDGB. Scale bar= 5 µm. **C.** Representative western blot showing RDGB can be co-immunoprecipitated along with dVAP-A from heads of 1 day old flies (N=3). **D.** Representative western blot showing RDGB^FFD/AAA^ cannot be co-immunoprecipitated along with dVAP-A from heads of 1 day old flies (N=2). **E.** Representative western blot showing RDGB^(USR1-LNS2)Δ^ cannot be co-immunoprecipitated along with dVAP-A from heads of 1 day old flies (N=3).

### An unstructured region of RDGB supports the interaction between FFAT/dVAP-A

The observation that RDGB^(USR1-LNS2)Δ^ could not engage in protein-protein interaction with dVAP-A suggests that additional regions C-terminal to the FFAT motif of RDGB are required to support this interaction. To identify this region, we made a series of progressively shorter deletions of RDGB starting from the C-terminus. To test the importance of the C-terminal LNS2 domain in supporting the FFAT/dVAP-A interaction, we deleted the C-terminal LNS2 domain from RDGB [Fig 1C-RDGB^LNS2Δ^] and expressed it in photoreceptors of *rdgB*^*9*^ flies (*rdgB*^*9*^; GMR>*rdgB*^LNS2Δ^) [Fig 3A]. We immunoprecipitated dVAP-A and found that RDGB^LNS2Δ^ could be detected in the dVAP-A pulled down fraction implying that the LNS2 domain is dispensable for FFAT/dVAP-A interaction [Fig 3B]. This finding suggests that the region between FFAT motif and the start of the LNS2 domain is essential for facilitating the FFAT/dVAP-A interaction.

**Figure 3:**
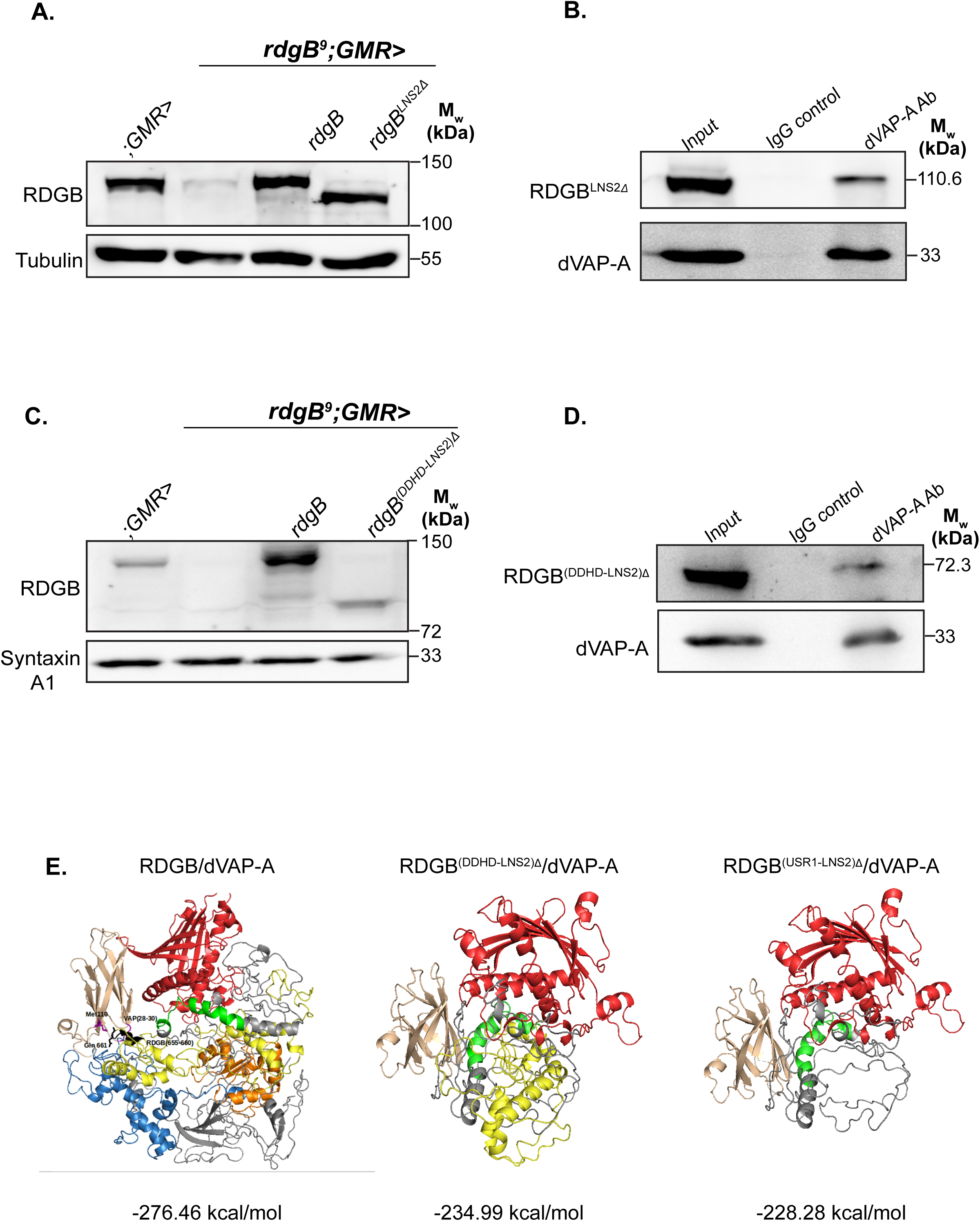
An unstructured region of RDGB supports the interaction between its FFAT motif and dVAP-A. A. Western blot of head extracts made from flies of the mentioned genotype. The blot is probed with antibody to RDGB. Tubulin is used as a loading control (N=3). B. Representative immunoblot showing RDGB^LNS2Δ^ can be co-immunoprecipitated along with dVAP-A from heads of 1 day old flies (N=3). C. Western blot of head extracts made from flies of the mentioned genotype. The blot is probed with antibody to RDGB. Syntaxin A1 is used as a loading control (N=3). D. Representative immunoblot showing RDGB^(DDHD-LNS2)Δ^ can be co-immunoprecipitated along with dVAP-A from heads of 1 day old flies (N=3). E. RDGB/dVAP-A interactions were analysed using protein-protein docking studies. The energies (Kcal/mol) for every complex (RDGB/dVAP-A, RDGB^(DDHD-LNS2)Δ^/dVAP-A, RDGB^(USR1-LNS2)Δ^/dVAP-A) have been marked below the representative structure. The PITPd protein is marked in red, FFAT motif in green, USR1 in yellow, the DDHD domain in blue and the LNS2 domain has been marked in orange. The dVAP-A protein is marked in pale orange.

The region between the FFAT motif and the LNS2 domain is mainly composed of a large unstructured region (USR1) and the DDHD domain. We tested the requirement of the DDHD domain in modulating the FFAT/dVAP-A interaction. Since LNS2 domain is dispensable for interaction with dVAP-A, we removed the C-terminal of the protein from just before the start of the DDHD domain [Fig 1C-RDGB^(DDHD-LNS2)Δ^] leaving the USR1 after the FFAT motif intact. We expressed this protein in fly photoreceptors (*rdgB*^*9*^; GMR>*rdgB*^(DDHD-LNS2)Δ^) [Fig 3C], and checked if it could interact with dVAP-A by immunoprecipitation. RDGB^(DDHD-LNS2)Δ^ was also found to coimmunoprecipitate with dVAP-A [Fig 3D]. Collectively, from the findings that RDGB^(DDHD-LNS2)Δ^ can interact with dVAP-A while RDGB^(USR1-LNS2)Δ^ cannot, suggest that the FFAT/dVAP-A interaction is supported by the unstructured region (USR1) present between the FFAT motif and the DDHD domain.

### Molecular docking analysis of the RDGB/dVAP-A interaction

To understand why an intact FFAT motif alone cannot bind dVAP-A, we performed protein-protein docking studies between dVAP-A and RDGB *in silico*. Since an experimentally determined structure is not available, an integrative modelling approach was used to obtain a 3-D structure of the full-length RDGB protein (details in materials and methods). Starting with this 3-D structure of wild type RDGB, we also derived 3-D structures of RDGB^(DDHD-LNS2)Δ^ and RDGB^(USR1-LNS2)Δ^. A structure for dVAP-A was obtained by homology modeling using the previously determined crystal structure of VAP-A (Kaiser et al., 2005). Each of the RDGB mutants was then used to perform docking studies with the structure of dVAP-A. Docking studies were performed using GRAMM-X (see materials and methods) and 20 models were generated for each of the three complexes (RDGB/dVAP-A, RDGB^(DDHD-LNS2)Δ^/dVAP-A and RDGB^(USR1-LNS2)Δ^/dVAP-A), and the energy and stability of each complex was calculated. For each of the three complexes, the best model (least energy) was selected and analyzed for interaction between the FFAT motif of RDGB protein and the MSP domain of dVAP-A. The energies of the complexes where we observed interactions between FFAT motif and MSP domain were in the order: RDGB/dVAP-A (−276.46 Kcal/mol) < RDGB^(DDHD-LNS2)Δ^/dVAP-A (−234.99 Kcal/mol) < RDGB^(USR1-LNS2)Δ^/dVAP-A (−228.28 Kcal/mol) [Fig 3E, **Supplementary Figure S1**]. This clearly indicates that the full-length RDGB/dVAP-A complex is most stable compared to RDGB^(DDHD-LNS2)Δ^/dVAP-A and RDGB^(USR1-LNS2)Δ^/dVAP-A complexes. Also the USR1 between the FFAT motif and DDHD domain [marked in orange-Fig 3E], was observed to interact with dVAP-A in the RDGB/dVAP-A and RDGB^(USR1-LNS2)Δ^/dVAP-A complex. This data thus provides a structural model to explain the requirement for the USR1 of RDGB in facilitating the interaction between dVAP-A and the FFAT motif of RDGB and supports the observation of loss of the interaction between RDGB and dVAP-A upon loss of USR1.

### An intact FFAT/dVAP-A interaction is not sufficient for proper localization of RDGB

We determined the sub-cellular localization of RDGB^(DDHD-LNS2)Δ^ and RDGB^LNS2Δ^, which can both interact with dVAP-A. Surprisingly, we found that neither RDGB^(DDHD-LNS2)Δ^ nor RDGB^LNS2Δ^ was localized to ER-PM contact sites as seen with RDGB [Fig 4A, B]. These results imply that while interaction with dVAP-A is essential, it is not sufficient for correct localization of RDGB to the ER-PM contact site at the base of the rhabdomere.

**Figure 4:**
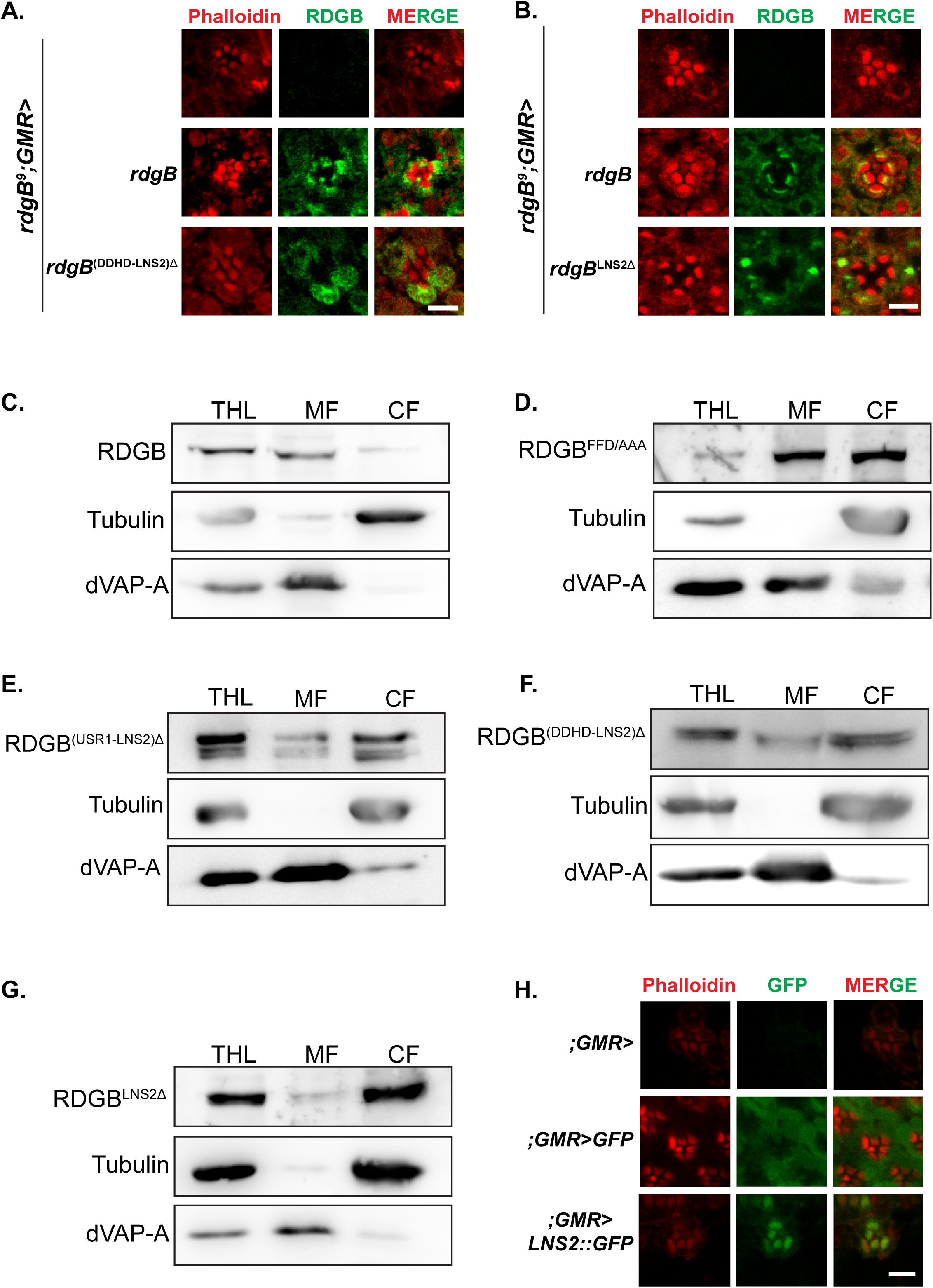
In addition to dVAP-A, apical plasma membrane association mediated by the LNS2 domain is indispensable for localization. A. Confocal images of retinae obtained from flies of the mentioned genotypes. Red represents phalloidin which marks the rhabdomeres and green represents RDGB. Scale bar= 5 µm. B. Confocal images of retinae obtained from flies of the mentioned genotypes. Red represents phalloidin which marks the rhabdomeres and green represents RDGB. Scale bar= 5 µm. C. Representative immunoblot showing RDGB cofractionates majorly in the membrane fraction from *Drosophila* heads. dVAP-A, an ER integral protein represents the membrane component, while the soluble protein tubulin represents the cytosolic pool [THL=Total Head Lysate, MF=Membrane fraction, CF=Cytosolic fraction] (N=3). D. Representative immunoblot showing RDGB^FFD/AAA^ cofractionates partially in the membrane fraction from *Drosophila* heads. dVAP-A, an ER integral protein represents the membrane component, while the soluble protein tubulin represents the cytosolic pool [THL=Total Head Lysate, MF=Membrane fraction, CF=Cytosolic fraction] (N=3). E. Representative immunoblot showing RDGB^(USR1-LNS2)Δ^ cofractionates majorly in the cytosolic pool from *Drosophila* heads. dVAP-A, an ER integral protein represents the membrane component, while the soluble protein tubulin represents the cytosolic pool [THL=Total Head Lysate, MF=Membrane fraction, CF=Cytosolic fraction] (N=3). F. Representative immunoblot showing RDGB^(DDHD-LNS2)Δ^ cofractionates majorly in the cytosolic pool from *Drosophila* heads. dVAP-A, an ER integral protein represents the membrane component, while the soluble protein tubulin represents the cytosolic pool [THL=Total Head Lysate, MF=Membrane fraction, CF=Cytosolic fraction] (N=3). G. Representative immunoblot showing RDGB^LNS2Δ^ cofractionates majorly in the cytosolic pool from *Drosophila* heads. dVAP-A, an ER integral protein represents the membrane component, while the soluble protein tubulin represents the cytosolic pool. [THL=Total Head Lysate, MF=Membrane fraction, CF=Cytosolic fraction] (N=3). H. Confocal images of retinae obtained from flies of the mentioned genotypes. Red represents phalloidin which marks the rhabdomeres and green represents GFP. Scale bar= 5 µm.

To test why the protein is mislocalized in spite of a stable interaction with dVAP-A, we performed a subcellular fractionation assay from *Drosophila* heads, and found that RDGB is a membrane associated protein that co-fractionates with the integral membrane marker dVAP-A [Fig 4C]. Surprisingly, mutation of the FFAT motif in RDGB (RDGB^FFD/AAA^), which disrupts the RDGB/dVAP-A interaction, led to only a partial loss of membrane association [Fig 4D]. These findings imply that factors other than an intact interaction with dVAP-A play a role in the membrane association and hence correct localization of RDGB.

### The LNS2 domain is essential for association of RDGB with the apical PM

We tested for any additional signals that may mediate the interaction of RDGB with membranes in *Drosophila* photoreceptors. Using the subcellular fractionation assay we found that RDGB^(USR1-LNS2)Δ^ majorly co-fractionated with the cytosolic marker tubulin [Fig 4E]. This finding suggests that the domains present C-terminal to the FFAT motif are required for membrane association. Upon sub-cellular fractionation of photoreceptors expressing RDGB^(DDHD-LNS2)Δ^, the majority of this protein cofractionates with the cytosolic marker tubulin suggesting that the membrane association domain is restricted to the DDHD and LNS2 domain [Fig 4F]. Finally, deletion of only the LNS2 domain from RDGB (RDGB^LNS2Δ^) led to a major loss in the membrane association of RDGB [Fig 4G]. These findings thus demonstrate that the LNS2 domain is indispensable for membrane association.

Having shown that the LNS2 domain is indispensable for interaction with the ER protein dVAP-A, we then sought to find the specific membrane in the photoreceptor that is bound by the domain. For this we took only the LNS2 domain of RDGB, tagged it to GFP and then expressed it in photoreceptors [**Supplementary Figure S2A**]. Unlike GFP which showed a completely diffused staining, LNS2::GFP was found to be localized specifically to the rhabdomeres, i.e. the apical PM [Fig 4H]. Upon expression in S2R+ cells, LNS2::GFP was also found to be localized to the PM **[Supplementary Figure S2B]**, further confirming that the LNS2 domain binds to PM in cells, and more specifically to the apical PM in photoreceptors which is part of the ER-PM MCS. Lipid binding assays revealed that the LNS2 domain binds phospholipids such as PA and PS **[Supplemental 2C]**, both of which are enriched at the PM. Thus, LNS2 in full length RDGB acts as an apical PM associating factor at ER-PM contact site by binding to PA and PS.

### Charged residues in the LNS2 domain of RDGB are required for membrane interaction and localization

We investigated the membrane binding mechanism of the LNS2 domain using molecular dynamics (MD) simulations (using the model generated as described in methods) and a dipalmitoylphosphatidylcholine (DPPC) membrane (system 1-**RDGB**). The system was subjected to six steps of equilibration at 50 ns per step followed by MD run for 100 ns (in replicates). The minimum distance between RDGB and the membrane in each system was measured throughout the 100 ns run to check the closest distance between RDGB and the DPPC membrane. Findings from this study show that the LNS2 domain is required for interaction of RDGB with apical PM. We captured the movements of the LNS2 domain during the simulation and calculated its distance from the membrane. As observed during the MD simulations, an alpha helical region of the LNS2 domain containing two charged residues was seen to move closest to the membrane. These were two lysine residues found at positions 1186 and 1187 of RDGB protein (K1186 and K1187).

The distance between K1186 and K1187 in the alpha helical region of the LNS2 domain and the membrane was then checked. It was observed that in the simulation with system 1-RDGB, the minimum distance between RDGB and the membrane is 2 Å which remains stable throughout the simulation **[Supplementary Figure S3A]**. The distance between the hydrogen atoms (closest to the lipid) of the residues K1186 and K1187 (LNS2 domain) and the DPPC molecule was 3 Å for upto 80% of the simulation after which it deviates to about 15 Å. Based on the above observation we mutated these two lysine residues to alanine (RDGB^KK/AA^) using the FOLDX protocol (referred in methods). This mutated protein was then subjected to similar MD simulations as for wild type RDGB (referred to as system 2-**RDGB^KK/AA^**). For system 2-RDGB^KK/AA^ the minimum distance between the protein and the membrane at the start of the simulation was 2 Å which increased to 8 Å as the simulation progressed **[Supplementary Figure S3B and S3C]**. The distance between the hydrogen atoms (closest to the lipid) of the residues A1186 and A1187 (in system 2-RDGB^KK/AA^) with DPPC molecule remained more than 5 Å for upto 60% of the simulation afterwhich it deviated to about 30 Å **[Supplementary Figure S3D,E and F]**. We also performed simulation for the protein lacking the LNS2 domain in the presence of membrane (system 3-**RDGB^LNS2Δ^**) and observed that the protein remained at a distance of 8 Å and more from the membrane throughout the simulation **[Supplementary Figure S3G]** and did not form any stable interactions. The charged residues of LNS2 domain in system 1-RDGB moved at a distance to the membrane such that they could form van der Waals interactions with each other while the distance between the protein and the membrane in system 2-RDGB^KK/AA^ and 3-RDGB^LNS2Δ^ remained larger throughout the simulation after the system stabilized **[Video files 1,2 and 3]**.

The study suggests that presence of LNS2 domain is important for interaction of the RBGB protein with the membrane. Since the minimum distance between the protein and the membrane is smaller in system 1(RDGB) as compared to system 2 (RDGB^KK/AA^) and system 3 (RDGB^LNS2[)5^), it is clear that this domain is required for the protein to associate to the membrane.

To validate the results of our simulations, we mutated the K1186 and K1187 residues of RDGB in the LNS2::GFP construct (LNS2^KK/AA^::GFP) and expressed it in S2R+ cells. Unlike wild type LNS2 which was specifically localized to the PM and some endomembrane structures, LNS2^KK/AA^::GFP was found to be present throughout the cell, with some residual binding to the PM also **[Supplementary Figure S3H]**. This observation is consistent with the observation in the MD simulation of system 2, where RDGB^KK/AA^ showed minimum interaction with the membrane at the start of the simulation and later deviated away from the membrane due to lack of stable interactions.

### The LNS2 domain is essential to support RDGB function during *in vivo* signalling

Given the indispensable role of LNS2 domain in localizing RDGB to the MCS, we tested if the domain has a physiological role in supporting RDGB function *in vivo*. A key function of RDGB is to maintain the electrical response to light in *Drosophila* photoreceptors (Harris and Stark, 1977; Yadav et al., 2015); this requirement manifests as a reduced electroretinogram (ERG) amplitude in *rdgB*^*9*^ flies. Upon testing the requirement of the LNS2 domain in supporting phototransduction, we found the light response in RDGB^LNS2Δ^ expressing photoreceptors to be as low as that in *rdgB*^*9*^ [Fig 5A, B]. RDGB is also essential to support the levels of PIP_2_ at the apical PM. We measured apical PM PIP_2_ levels by quantifying the fluorescence of PH-PLCδ::GFP probe in the pseudopupil of the eye (Chakrabarti et al., 2015). As previously reported (Yadav et al., 2015),we found that the resting level of PIP_2_ at the apical PM of *rdgB^9^* was reduced and could be restored to wild type levels by reconstitution with a wild type RDGB transgene [Fig 5C, D]. However, when *rdgB*^*9*^ was reconstituted with RDGB^LNS2Δ^ (*rdgB*^*9*^; *GMR> rdgB*^LNS2Δ^), the PM PIP_2_ levels were found to be as low as in *rdgB*^*9*^ photoreceptors [Fig 5C, D], despite equivalent levels of probe expression [Fig 5E]. These results collectively support the indispensable role of the LNS2 domain in supporting RDGB function *in vivo*.

**Figure 5:**
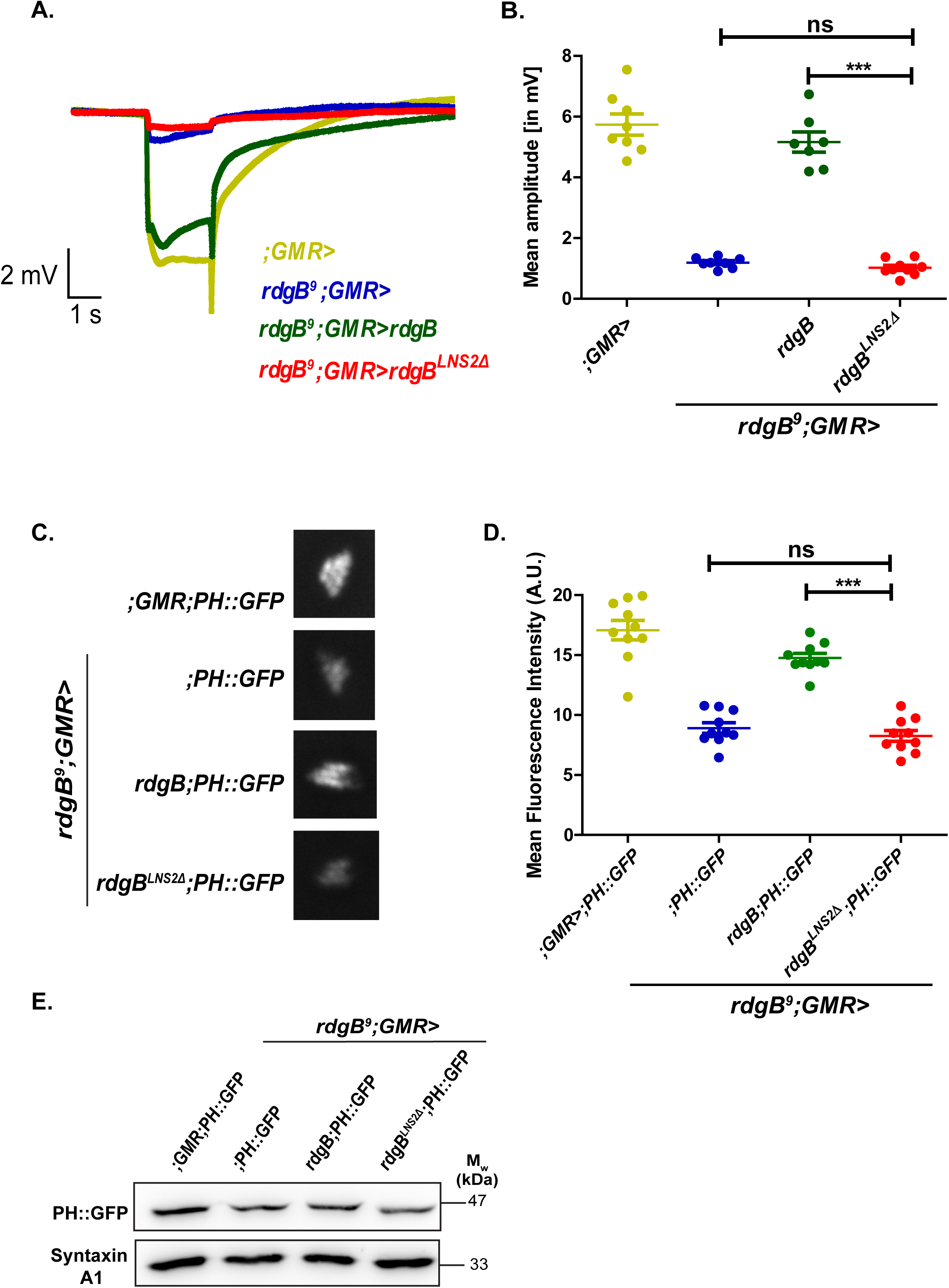
The LNS_2_ domain is indispensable for supporting RDGB function *in vivo*. A. Representative ERG trace of 1 day old flies of the mentioned genotypes. Y-axis represents amplitude in mV, X-axis represents time in sec. B. Quantification of the light response from 1 day old flies of the mentioned genotype. Each point on Y-axis represents mean amplitude ±s.e.m., X-axis represents genotype (*** - p<0.001, ns= not significant; two tailed unpaired t-test). C. Representative images of fluorescent deep pseudopupil from 1 day old flies of the mentioned genotypes expressing the PH-PLCδ::GFP probe. D. Quantification of the fluorescence intensity of the deep pseudopupil (A.U. =Arbitrary Units). Y-axis denotes the mean intensity per unit area ±s.e.m., X-axis denotes genotype. (*** - p<0.001, two tailed unpaired t-test) E. Western blot of head extracts made from flies of the mentioned genotypes expressing PH-PLCδ::GFP probe. The blot is probed with antibody to GFP. SyntaxinA1 is used as a loading control (N=3).

### The LNS2 domain restricts the movement of the DDHD domain

While the data above highlights the indispensable role of LNS2 domain in supporting RDGB function *in vivo*, previous studies (Milligan et al., 1997; Yadav et al., 2015) have reported the sufficiency of the PITPd in supporting RDGB function *in vivo*. We obtained similar results when we tried to rescue *rdgB*^*9*^ phenotypes by expressing RDGB^(USR1-LNS2)Δ^. RDGB^(USR1-LNS2)Δ^ was able to completely rescue the reduced light response phenotype of *rdgB*^*9*^ mutants [Fig 6A, B]. Similarly, expression of RDGB^(USR1-LNS2)Δ^ could partially restore the reduced basal PIP_2_levels in *rdgB*^*9*^ photoreceptors to wild type levels [Fig 6C,D], with equivalent levels of PH-PLCδ::GFP probe expression [Fig 6E]. Importantly, while loss of the LNS2 domain completely abrogated *in vivo* RDGB function, loss of all the domains collectively after the PITPd has only marginal effect on its function.

**Figure 6:**
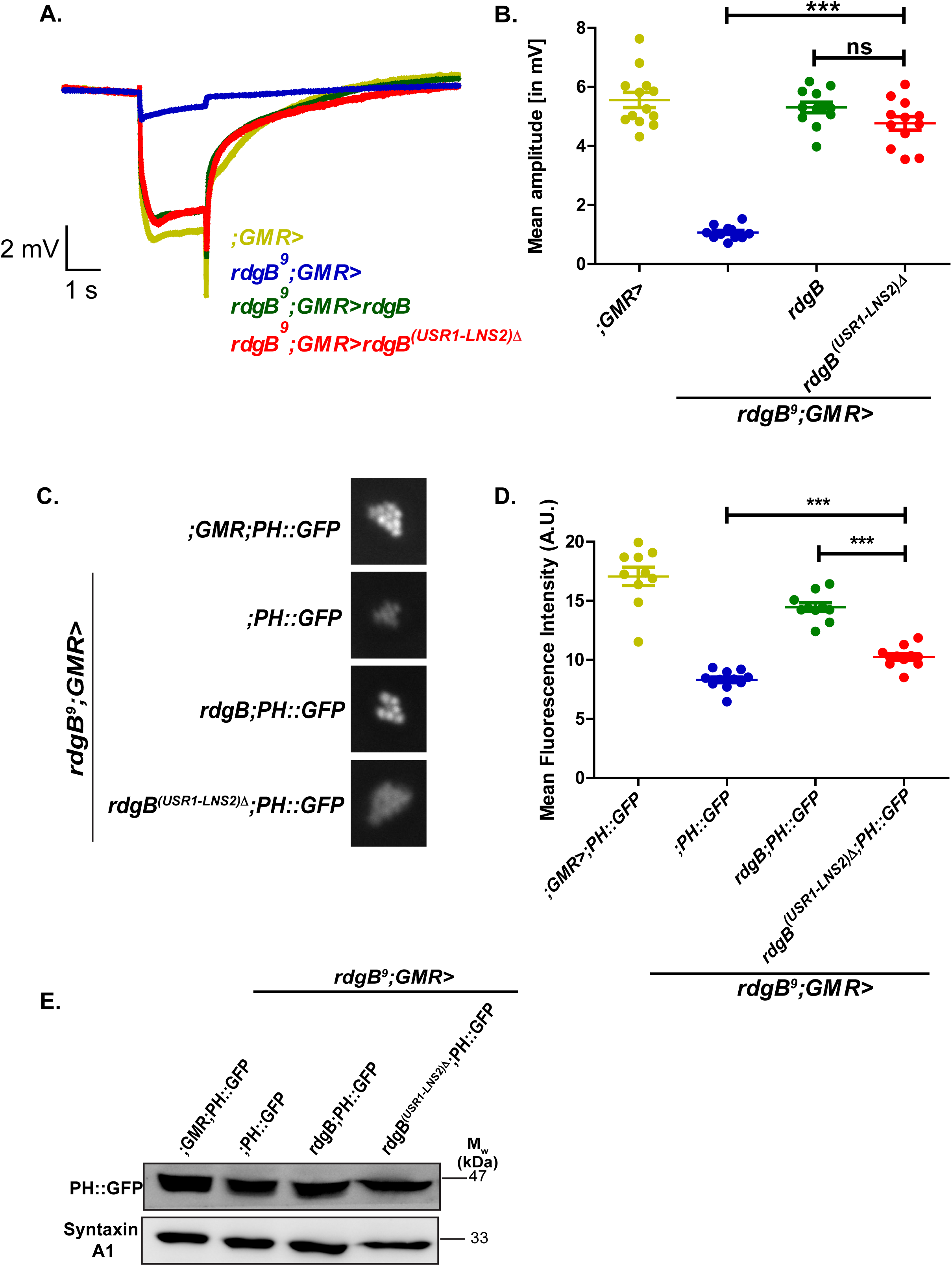
Loss of all the C-terminal domains only partially affect RDGB function. A. Representative ERG trace of 1 day old flies of the mentioned genotypes. Y-axis represents amplitude in mV, X-axis represents time in sec. B. Quantification of the light response from 1 day old flies of the mentioned genotype. Each point on Y-axis represents mean amplitude ±s.e.m., X-axis represents genotype (*** - p<0.001, ns= not significant; two tailed unpaired t-test). C. Representative images of fluorescent deep pseudopupil from 1 day old flies of the following genotypes expressing the PH-PLCδ::GFP probe. D. Quantification of the fluorescence intensity of the deep pseudopupil (A.U. =Arbitrary Units). Y-axis denotes the mean intensity per unit area ±s.e.m., X-axis denotes genotype (*** - p<0.001, two tailed unpaired t-test). E. Western blot of head extracts made of flies of the mentioned genotypes, expressing PH-PLCδ::GFP probe. The blot is probed with antibody to GFP. SyntaxinA1 is used as a loading control (N=3).

One possible explanation for this conundrum is that the C-terminal domains might engage in interdomain interaction in the context of the full length protein. To understand the interdomain interactions of C-terminal domains (DDHD and LNS2) with the PITPd, truncated protein models were derived from the full-length model. Two models were generated, one without the LNS2 domain (RDGB^LNS2Δ^) and the other without the USR1, DDHD and LNS2 domains[RDGB^(USR1-LNS2)Δ^]. In order to predict potential inter-domain interactions, a normal mode analysis (described in materials and methods) was performed. Lowest frequency modes that describe the largest movements were analysed in detail for all the three models.

In all the low frequency modes of full-length RDGB protein, the DDHD domain moved towards the LNS2 domain while the PITPd show limited interaction with the other domains. This most favourable mode indicates that there exist inter-domain interactions between the two C-terminal domains (DDHD and LNS2) of RDGB [Figure 7(A) and **Video file 4]**. There being no large structural changes or movements in the PITPd of RDGB across domains, the function of the protein remains conserved as seen in data mentioned above.

**Figure 7:**
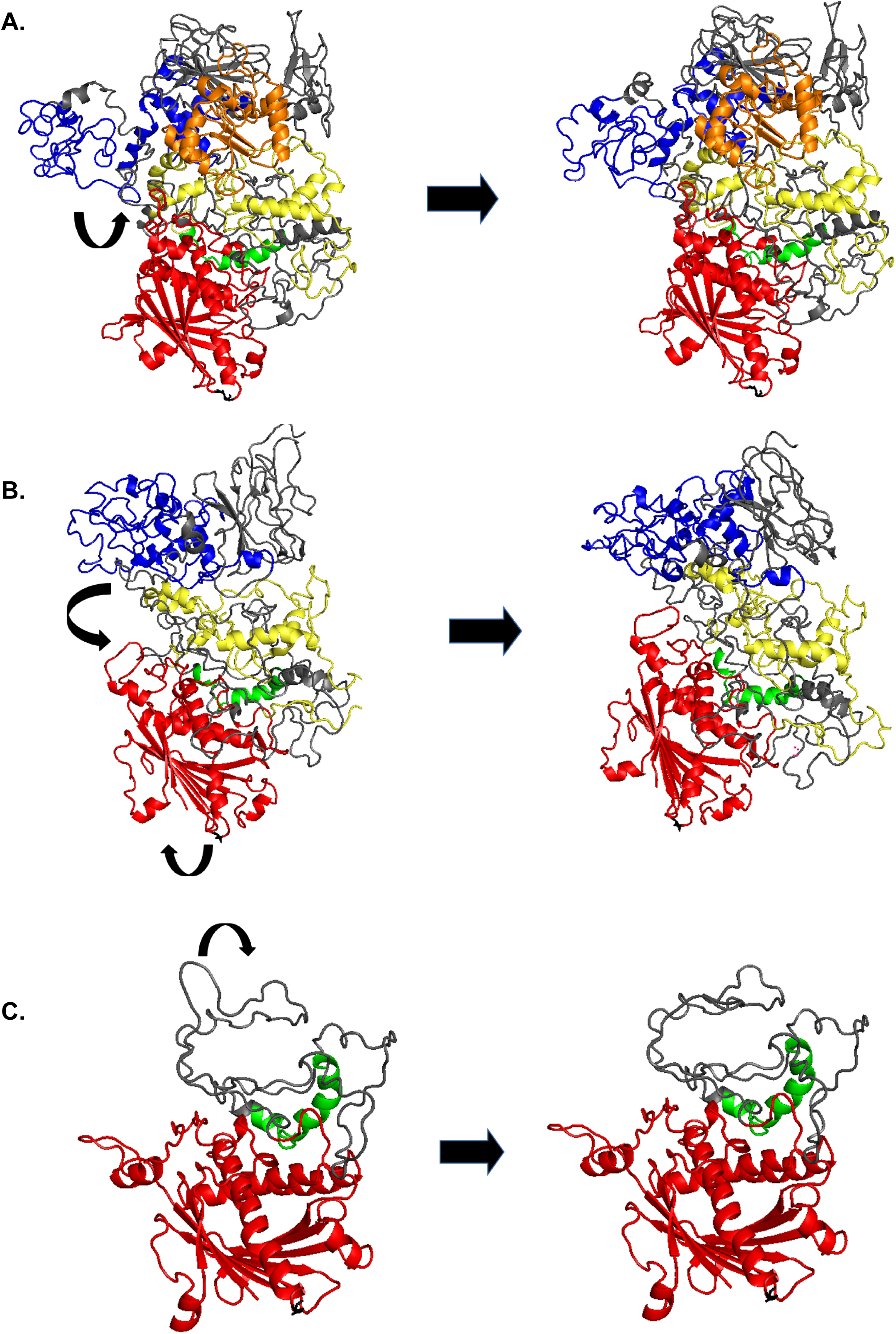
Normal model analysis of RDGB, RDGB^LNS2Δ^ and RDGB^(USR1-LNS2)Δ^ protein: Normal mode analysis was carried for RDGB protein and its C-terminal deletion variants to understand inter-domain movements leading to domain interaction. PITPd is marked in red, FFAT motif in green, USR1 in yellow, DDHD domain in blue and LNS2 domain in orange. The ‘YW’ amino acid pair important for membrane docking is marked in black. The arrows indicate the direction and amplitude of domain movements. A. RDGB, B. RDGB^LNS2Δ^ and C. RDGB^(USR1-LNS2)Δ^.

In contrast, the normal mode movements of RDGB^LNS2 Δ^ revealed a large conformational change in the protein in the absence of LNS2 domain. In the protein model of RDGB^LNS2Δ^, it was observed that the DDHD domain folded back on the PITPd [Figure 7(B) and **Video file 5]**. This movement also resulted in bending of the PITPd unlike the lack of movement of this domain observed in the full-length RDGB. Although this movement does not affect the lipid binding residues of PITPd, there is a large movement seen for the YW motif as compared to wild type. The WW motif present in mammalian PITPα and PITPβ is essential for membrane docking (Shadan et al., 2008) and mutation of the equivalent YW motif in the PITPd of full-length RDGB leads to complete loss of RDGB function (Yadav et al., 2015). Since membrane docking is the primary step for lipid transfer by PITPs, it is possible that deletion of the LNS2 domain affects membrane docking of the PITPd resulting in complete loss of RDGB function.

By contrast, in RDGB^(USR1-LNS2)Δ^, all the 10 modes showed no movement of the PITPd and FFAT motif. The only movement detected was of the two loops containing 6 residues each at the edges of FFAT motif [Figure 7(C) and **Video file 6]** and the movement of YW motif was unaffected upon deletion of the entire C-terminus. Since the RDGB^(USR1-LNS2)Δ^ protein was able to function normally, these findings strongly suggest that an intact PITPd with limited or no inter-domain movements is sufficient for normal functioning of RDGB protein.

## Discussion

The presence of multiple domains in LTPs is hypothesized to enable their correct localization at MCS. These domains are conceptualized as independent units each with a unique property contributing to optimal lipid transfer function at MCS. A similar model has been proposed for a specific group of LTPs, the PITPs that transfer PI at ER-PM junctions (Kim et al., 2015; Kim et al., 2013). However, in the case of *Drosophila* RDGB, a multidomain PITP, it has been noted that re-expression of just the PITPd of RDGB (RDGB^PITPd^) which performs lipid transfer *in vitro*, in a null mutant background, is sufficient to rescue key phenotypes *in vivo* (Milligan et al., 1997; Yadav et al., 2015) suggesting the sufficiency of the RDGB^PITPd^ in supporting RDGB function. A more recent study has however shown that while RDGB^PITPd^ can rescue key phenotypes, it is incapable of supporting lipid turn over during high rates of PLC-β signalling (Yadav et al., 2018), emphasizing the importance of ensuring a sufficiently high concentration of RDGB at the ER-PM contact site in photoreceptors.

How is RDGB accurately localized to ER-PM MCS? It has previously been reported (Yadav et al., 2018) that an interaction between the FFAT motif and the ER-resident protein dVAP-A is essential for the normal localization and function of RDGB. In this study, surprisingly, we found that a RDGB protein lacking all domains downstream of the FFAT motif was (i) mislocalized away from the base of the rhabdomere and (ii) was unable to interact with dVAP-A despite the presence of an intact FFAT motif, suggesting that additional regions of the RDGB protein are required to stabilize the FFAT/dVAP-A interaction in this protein. Using a series of deletion constructs of RDGB, and MD simulations *in silico*, we established that the unstructured region 1 (USR1), positioned C-terminal to the FFAT motif but prior to the start of the DDHD domain is sufficient to stabilize the FFAT/dVAP-A interaction. These findings suggest that inter-domain interactions within the RDGB protein are important for stabilizing the interaction between its FFAT motif and dVAP-A, thus anchoring it to the ER side of the ER-PM contact site. Interestingly, a role for an intrinsically disordered region of OSBP in regulating its function at ER-Golgi MCS has recently been proposed (Jamecna et al., 2019). In the broader context, a large number of proteins that interact with dVAP-A have been reported (Murphy and Levine, 2016); many of these are via interaction with the FFAT motif but similar to the findings reported in this manuscript, they too may require other domains to stabilize this interaction. This may offer additional modes of recruiting regulators of lipid transfer proteins to the vicinity of the contact site.

Proteins localized to an ER-PM MCS also require a mechanism to tether them to the PM. In this study, using sub-cellular fractionation assay, we found that RDGB is a membrane associated protein. This membrane association is likely to be both with the ER and the apical PM of photoreceptors. Disruption of RDGB/dVAP-A interaction only reduced membrane association of RDGB partially, implying the presence of additional signals that allow RDGB to associate with membranes. Using a series of C-terminal deletion constructs, we found that this key membrane association signal is the C-terminal LNS2 domain. When the LNS2 domain is deleted (RDGB^LNS2Δ^), the majority of the protein is cytosolic and is mis-localized away from the ER-PM contact site into the photoreceptor cell body. Upon expression of the LNS2 domain alone, it was found to be targeted to the apical PM in photoreceptors. Lipid binding assay revealed that the LNS2 domain is able to bind PM enriched acidic phospholipids such as PA and PS *in vitro*, suggesting a molecular signal from the PM that engages in this interaction. The contribution of these signals to function *in vivo* is clearly critical since RDGB^LNS2Δ^ is unable to restore function when reconstituted in the *rdgB*^*9*^ mutant. Interestingly, the protein-protein interaction of RDGB^LNS2Δ^ with dVAP-A itself is unaffected, implying that the interaction with dVAP-A and apical PM, are mutually exclusive properties of RDGB. These findings suggest that the function of the LNS2 domain is a prerequisite to target RDGB to the apical PM, and following this the USR1 assisted FFAT/dVAP-A interaction restricts its localization specifically to the apical PM-ER junction. Based on these findings, the RDGB protein can be regarded as a PITP localized to ER-PM MCS sites through the interaction of its FFAT domain and USR1 on the ER side of the MCS and through its LNS2 domain on the PM side **[Graphical Abstract]**.

Although we demonstrate key roles for the FFAT motif, USR1 and the LNS2 domains, in RDGB function, why is the PITPd able to rescue key phenotypes when expressed at adequate levels on its own (Milligan et al., 1997; Yadav et al., 2015)? We found that while deletion of the LNS2 domain alone led to complete loss of RDGB function, loss of USR1, DDHD and LNS2 domains together only affect RDGB function partially. Through protein structure modelling of RDGB, we found that in the full-length protein, the movement of the PITPd movement is restricted while the C-terminal domains interact, presumably allowing the PITPd to shuttle lipids between the ER and PM. Loss of the LNS2 domain alone results in the DDHD domain folding back to the PITPd. The folding back of the DDHD domain on the PITPd restricts the movement of the membrane docking YW motif, thus inhibiting RDGB function *in vivo* as a whole **[Graphical Abstract]**. These observations underscore the molecular choreography between the movements of individual domains of the RDGB protein that are critical to deliver its function. Interestingly, a bioinformatics analysis of all available genome sequences for the domain composition of proteins containing a PITPd reveals that in nature proteins exist in a number of varieties in which the PITPd is present in conjunction with additional domains. These additional domains may include a DDHD domain alone or in conjunction with further domains such as but not restricted to the LNS2 domain. Such combinations of domains may have evolved to accurately localize the PITPd to specific parts of the cell or to regulate its function.

In summary, our study identifies two novel intramolecular interactions that are required to facilitate accurate localization of RDGB at ER-PM contact sites. In addition, this study provides evidence for interdomain interactions in choreographing the molecular dynamics of the proteins contributing to its lipid transfer activity at this MCS. In the absence of either of these signals the RDGB protein is mislocalized away from the ER-PM contact site with profound consequences for PM PIP_2_ level homeostasis during PLC-β signalling. Our findings also provide a conceptual framework for understanding the regulation of PI transfer activity at MCS.

## Supporting information

Basak_Video1

Basak_Video2

Basak_Video3

Basak_Video4

Basak_Video5

Basak_Video6

## Acknowledgements

This work was supported by the National Centre for Biological Sciences-TIFR and a Wellcome-DBT India Alliance Senior Fellowship (IA/S/14/2/501540) to PR. We thank the transgenic fly facility, Central Imaging Facility and High-Performance Computing Facility at NCBS for support. We would also like to thank Dr. Girish Ratnaparkhi from IISER Pune for providing us with dVAP-A antibody.

## Materials and methods

### Fly stocks

All fly stocks were maintained at 25°C incubators with no internal illumination. Flies were raised on standard corn meal media containing 1.5% yeast. UAS-Gal4 system was used to drive expression in the transgenic flies.

### Molecular Biology

BDGP gold clone 09970 containing the *rdgB*-RA transcript was used as the parent vector for making various constructs of RDGB used for the experiments. The cDNA coding region corresponding to RDGB^(USR1-LNS2)Δ^ (amino acids 1-472) was subcloned into pUAST-attB by using the restriction enzymes *NotI* and *XbaI* (NEB). Similarly, for making RDGB^(DDHD-LNS2)Δ^ the cDNA corresponding to amino acids 1-655 was amplified, and for RDGB^LNS2Δ^ the cDNA corresponding to amino acids 1-1000 was amplified and then individually subcloned in *NotI* and *XbaI* digested pUAST-attB. For cloning of the LNS2 domain, the cDNA of RDGB corresponding to amino acids 947-1259 was subcloned in pJFRC::GFP vector using the restriction enzymes *BglII* and *NotI* (NEB). A flexible linker of Gly(G)-Ser(S) of the sequence G-G-S-G-G-G-S-G-G-G-S-G-G was introduced between the LNS2 domain and GFP to allow independent and efficient folding of the two proteins. For cloning of LNS2^KK/AA^, site directed mutagenesis was done in the pJFRC-LNS2::GFP clone. Lysine residues corresponding to 1184 and 1185 positions in RDGB were mutated to alanine in pJFRC-LNS2::GFP by introducing the mutations directly in the primers used for amplification.

### Cell culture, transfection and immunofluorescence

S2R+ cells were cultured in Schneider’s insect medium (HiMedia) supplemented with 10% Fetal Bovine Serum and with antibiotics Penicillin and Streptomycin. Cells were transfected using Effectene (Qiagen) as per manufacturer’s protocol. Post 24 hours of transfection, cells were fixed with 4% paraformaldehyde (Electron Microscopy Sciences) and imaged to observe for GFP fluorescence using a 60X 1.4 NA objective, in Olympus FV 3000 microscope

### Western Blotting

Heads of one day old flies were homogenised in 2X Laemmli sample buffer, and boiled at 95°C for 5 minutes. The samples were then run on a SDS-PAGE gel, and transferred on to a nitrocellulose membrane [Hybond-C Extra; (GE Healthcare, Buckinghamshire, UK)], with the help of a semi-dry transfer apparatus (BioRad, California, USA). The membrane was then blocked using 5% Blotto (sc-2325, Santa Cruz Biotechnology, Texas, USA) in Phosphate-buffered saline (PBS) with 0.1% Tween 20 (Sigma Aldrich) (PBST) for 2 hrs at room temperature (RT). The membrane was then incubated with the respective primary antibody, overnight at 4°C, using the appropriate dilutions [anti-RDGB (lab generated), 1:4000; anti-dVAP-A (kind gift from Dr. Girish Ratnaparkhi, IISER Pune), 1:3000; anti-α-tubulin-E7 (DSHB, Iowa, USA), 1:4000; anti-syntaxinA-8C3 (DSHB, Iowa, USA), 1:1000; anti-GFP (sc-9996), 1:2000]. Following this, the membrane was washed in PBST, and incubated with the appropriate secondary antibody (Jackson Immunochemicals; dilution used: 1:10,000) coupled to horseradish peroxidase, at RT for 2 hrs. The blots were visualized using ECL (GE Healthcare), and imaged in a LAS4000 instrument.

### Immunostaining

For immunohistochemistry, retinae of one-day old flies were dissected under bright light in PBS. The samples were then fixed using 4% paraformaldehyde (Electron Microscopy Sciences) in PBS with 1 mg/ml saponin (Sigma Aldrich) for 30 minutes at RT. Post fixation, samples were washed thrice with PBS having 0.3% Triton X-100 (PBTX) and blocked using 5% Fetal Bovine Serum (ThermoFisher Scientific) in PBTX for 2 hrs at RT. The samples were then incubated overnight with the appropriate antibody in blocking solution at 4°C [anti-RDGB, (1:300); anti-GFP (1:5000), ab13970 (Abcam Cambridge, UK)]. Samples were then washed thrice with PBTX and incubated with the secondary antibody [Alexa Fluor 633 anti-rat (A21094), anti-chick (A21103), IgG (Molecular Probes)] at 1:300 dilution for 4 hrs at RT. For staining of the F-actin, Alexa Fluor 568–Phalloidin (Invitrogen, A12380) at 1:300 dilution was added during incubation with the secondary antibody. Samples were then washed in PBTX thrice and mounted with 70% glycerol in PBS. The whole-mounted preparations were imaged under 60X 1.4 NA objective, in Olympus FV 3000 microscope.

### Co-immunoprecipitation

Snap-frozen *Drosophila* heads were lysed in ice-cold Protein Lysis Buffer [50mM Tris-Cl, 1mM EGTA, 1mM EDTA, 1% Triton X-100, 50mM NaF, 0.27 M Sucrose, 0.1% β-Mercaptoethanol]. 10% of the head lysate was aliquoted to be used as input. The remaining lysate was split into two equal parts. To one part, 2 µl of dVAP-A antibody was added, and to the other part, 2 µl of control IgG was added, and incubated overnight at 4°C. On the next day, Protein-G sepharose beads (GE Healthcare) were spun at 13000X g for 1 minute, and then washed with Tris-buffered saline (TBS), twice. The beads were then incubated with 5% Bovine Serum Albumin (BSA) (HiMedia) in TBS with 0.1%Tween-20 (TBST) for 2 hrs at 4°C. Equal amounts of blocked beads were then added to each sample, and incubated at 4°C for another 2 hrs. The immunoprecipitates were then washed twice with TBST containing 2-Mercaptoethanol, and 0.1 mM EGTA for 5 minutes. The supernatant was then removed, and the beads were boiled in 2X Laemmli sample buffer for western blotting.

### Sub-cellular fractionation assay

The assay was performed as described by Sanxaridis et al., 2007 with minor modifications. Briefly, snap-frozen *Drosophila* heads were homogenised in ice-cold homogenisation buffer A (30 mM NaCl, 20 mM HEPES, 5 mM EDTA, pH=7.5). 10% of homogenate, representing the total head lysate, was directly taken for western blotting. The remaining homogenate was centrifuged at 5000 rpm for 5 minutes at 4°C to remove all chitinous material. The pellet was re-homogenized in the buffer to redeem any remaining membranous component from the cell ghost. This was done twice, post which the homogenate was spun at 100,000X g for 30 minutes, at 4°C to separate the entire membranous component from the cytosolic fraction. The pellet was reconstituted in buffer A. The re-suspended pellet representing the membrane fraction, and the supernatant representing the cytosolic fraction, were then individually used for doing western blotting.

### Electrophysiology

Anaesthetised flies were immobilized at the end of a pipette tip by applying a drop of colourless nail polish on the proboscis. For recordings, GC 100F-10 borosilicate glass capillaries (640786, Harvard Apparatus, MA) were pulled to form electrodes and then filled with 0.8% (w/v) NaCl. The reference electrode was placed on the centre of the eye and the ground electrode on the thorax to obtain voltage changes post stimulation. The protocol for recording involved dark adapting the flies for 5 minutes initially, following which they were shown green flashes of light for 2 secs (10 times), and 12 secs of recovery time in dark between the two flashes. Voltage changes obtained were amplified using DAM50 amplifier (SYS-DAM50, WPI, FL), and recorded using pCLAMP10.7. Analysis was done using Clampfit 10.7 (Molecular Devices, CA). For analysis, the average of 10 recordings was taken for per fly.

### Deep pseudopupil imaging

The imaging is done with flies expressing a single copy of PH-PLCδ::GFP (PH domain of PLCδ, a PIP_2_biosensor, tagged to GFP) driven by the transient receptor (*trp*) promoter of flies. Flies were anaesthetised and immobilized at the end of a pipette tip using a drop of colourless nail polish. The flies were then placed on the stage of an Olympus IX71 microscope, and the fluorescent pseudopupil focussed using a 10X objective lens. For imaging the deep pseudopupil (DPP), the flies were first adapted to red light for 6 minutes, following which a blue flash of 90 msec was given. The emitted fluorescence was captured, and its intensity was measured using Image J from NIH (Bethesda, Maryland, USA). Quantification of the DPP fluorescence intensity was done by measuring the intensity values per unit area of the pseudopupil. The values are represented as mean +/− s.e.m.

### Lipid overlay assay

S2R+ cells were transfected with pJFRC-LNS2::GFP and pJRFC::GFP for 48 hours, following which cells were lysed with Protein Lysis Buffer (50 mM Tris-Cl, 1 mM EGTA, 1 mM EDTA, 1% Triton X-100, 50 mM NaF, 0.27 M Sucrose, 0.1% β-Mercaptoethanol). 10% of the cell lysate was aliquoted separately to be used for western blotting. In parallel, commercially available PIP strips (Echelon Biosciences, P-6001) were blocked using 5% BSA (HiMedia) in TBST for 2 hours at RT. Following this, the strips were incubated overnight at 4°C with the remaining cell lysate. Next the membranes were washed extensively 5 times with 0.1% TBST and then incubated with anti-GFP antibody [(sc-9996), 1:2000] at RT for 2 hours. The membranes were then probed with HRP-conjugated anti-mouse IgG (Jackson Immunochemicals; 1:10,000) and binding was detected using ECL (GE Healthcare) in a LAS4000 instrument.

### Structure prediction of RDGB

*In-silico* integrative modelling was done to predict RDGB structure. RDGB is a multi-domain protein and contains the following domains-PITPd (1-280 aa), FFAT-motif (396-416 aa), DDHD domain (711-900 aa) and LNS2 domain (1159-1200 aa). There is an unstructured region-USR1 (no secondary structure predicted) between the FFAT-motif and DDHD domain (420-600 aa). Secondary structure prediction shows that the PITPd contains alpha helices and beta sheets, the DDHD domain is mostly alpha helical and the LNS2 domain contains alpha helices and beta sheets. The region between the PITPd and DDHD domain is largely unstructured and lacks high homology to any existing protein structure. In order to obtain a 3-D structure of the protein, homology-based modelling (MOELLER) (Roy et al., 2010) was done [Table 1]. The regions which have high homology to known structures available in PDB database were modelled using the homology modelling protocol in MODELLER. The regions that did not have template predicted with high identity but had closely related protein FOLD (classified as per SCOP classification) structure were modelled using fold prediction method while those regions of the model which had neither of the two above mentioned templates were modelled *ab-initio*. The table lists all templates that were obtained by either homology or fold prediction, while regions where template coverage was low were modelled using *ab-initio* methods. The final full-length model was energy minimized for 500 steps of conjugate gradient cycles followed by steepest descent gradient for 500 steps. The initial steps of minimization were stringent to avoid large structural deviations in the model while minimization. Once the model reached a negative potential energy state with allowed Ramachandran plot values, relaxed energy minimization steps were performed to further reach an energy minimum. The final model had negative potential energy and is stable. The Ramachandran plot distribution of amino acids in the model was checked using the PROCHECK analysis tool (Laskowski et al., 1993). To further check the stability of the protein in solution, a molecular dynamics simulation was carried out for 100 ns using the DESMOND module. The protein RMSD reached 7 Å at 20 ns and remained stable until 100 ns. Similarly, the RMSF of the protein at regions of stable secondary structure remained below 6 Å throughout the simulation except for the unstructured regions which show a higher RMSF.

**Table 1:** Details on modelling protocol for RDGB protein. The table represents the details on templates used for modelling the RDGB protein using an integrative modelling approach. The first column represents the different regions of RDGB protein modelled individually using different templates. The second and third column represents the available template and the protein name of the template. The fourth and fifth column represent the confidence (in percentage) with which the template is selected for modelling and percent identity of the template with the RDGB protein sequence respectively. The last column indicates the method used for modelling. The well-structured domains of the RDGB protein were modelled using the available templates, either by homology modelling or fold prediction. For regions with low confidence or low sequence identity with the template, *ab-initio* modelling protocol was used.

| RDGB sequence | Template | Protein name | Confidence | % Identity | Method used for modelling |
| --- | --- | --- | --- | --- | --- |
| **1-300 (PITP)** | 1t27a | PITP | 100 | 44 | Homology modelling |
|  | 1kcma | PITP | 100 | 52 |  |
| **300-420 (FFAT)** | 2rr3 | Complex b/w VAP-A and human OSBPFFAT motif protein |  |  | Hhpred(distant homology) |
| **420-600** | 1auaa | Phospholipid binding protein sec14p | 69.6 | 13 | Fold prediction |
| **600-700** | 3uuea | Hydrolase-lip1secretory lipase | 95.5 | 15 | Fold prediction |
| **700-800 (DHDD)** | 1dwka | Cyanase C-terminal domain | 16.5 | 4.6 | Loop modelling and fold prediction |
|  | 3ebca | Hydrolase | 12.6 | 30 | Loop modelling and fold prediction |
| **800-900 (DHDD)** | 2pqrd | WD repeat protein | 21.7 | 63 | Modelled ab initio/templatecoverage low |
|  | 4z38b | Transferase/enoyl reductase | 13.7 | 18 | Modelled ab initio/templatecoverage low |
| **900-1059** | 1f86a | Transthyretin protein | 97 | 21 | Fold prediction |
| **1059-1200 (LNS-2)** | 1ltqa/1xpja | HAD like phosphatase domain of polynucleotide kinase | 99.9/99.8 | 13 | Homology modelling |
| **1200-1284** | 5d5pc | transferase | 14-17% | 26-36% | 58% disordered region –ab initio modelling |

### Molecular docking of RDGB, RDGB^(USR1-LNS2)Δ^ and RDGB^LNS2Δ^ to dVAP-A

Molecular docking was performed for each of the RDGB models with that of dVAP-A [obtained by homology modeling using dVAP-A X-ray crystal structure (from *Rattus novergicus*)]. GRAMM-X protocol (Tovchigrechko and Vakser, 2005) was used to carry out the docking studies. GRAMM-X follows the Fast Fourier Transformation (FFT) methodology. It uses a smoothed Lennard-Jones potential on a fine grid during the global search FFT stage, followed by the refinement optimization in continuous coordinates and then performs re-scoring of the complexes using the knowledge-based potential terms for protein-protein interactions. 20 models were generated for every set of protein-protein interactions. The energy values were calculated for all the 60 models using the PPcheck algorithm (Sukhwal and Sowdhamini, 2013). The models with low potential energy values were selected to understand the interaction of RDGB with dVAP-A protein. Each of the low energy complex was manually visualized in PYMOL (The PyMOLMolecular Graphics System, Version 2.0 Schrödinger, LLC.) for interactions between FFAT motif of the RDGB protein and MSP domain of dVAP-A. The models which had the relevant known interactions between the two proteins were selected for further analysis. The protein-protein complex was analyzed for stable interactions based on PPcheck evaluation and models with lowest overall energy and very low unfavorable interaction was selected as the best model. Each of the best models selected was analyzed further using molecular dynamics simulation for stability of the complex in solution. Desmond module of molecular dynamics was used to perform 100 ns simulation for each protein in duplicates. OPLS_2005 force field with standard NPT conditions was used. The protein complexes were solvated in an orthorhombic box with periodic boundary conditions by adding TIP3P water molecules. The initial equilibration was carried out using default protocol of restrained minimization followed by molecular dynamics simulations for 100 ns. Based on the analysis described below we obtained a RDGB model with K1186 and K1187 residues mutated to alanine using FOLDX (Schymkowitz et al., 2005). The repair model and build model module of FOLDX was used along with rotamer stabilization to obtain the structure of the mutant protein. The mutant protein was as stable as the wild type as measured using FOLDX energy values.

### Molecular dynamics (MD) simulation of RDGB interactions at a membrane

MD simulations were done with full length RDGB protein in the presence of a DPPC membrane to identify the domains and residues important for interaction of the protein with the membrane. CHARMM-GUI membrane builder module (http://www.charmm-gui.org/?doc=input/membrane.bilayer) was used to generate the membrane input file for Gromacs MD simulations. A DPPC membrane with 250 molecules of lipids each on upper and lower leaflet of the membrane was built using the CHARMM-GUI membrane builder. Three independent membrane-protein systems were generated for RDGB protein (system 1), RDGB^KK/AA^ (system 2) and the RDGB^LNS2Δ^ protein (system 3). The protein was placed at a distance of 10 nm along the Z-axis from the membrane at the start of the simulation. The protein and the membrane system were solvated in an orthogonal box with TIP3P water and neutralized with ions to get a final system with net charge being zero. The simulations were performed with a 2 fs step size and the nearest neighbour list was recorded every 20 ps. The temperature and pressure were maintained at 300K and 1 bar^−1^ respectively. The system was energy minimized using Gromacs charmm36m force field for 50 ns followed by six steps of equilibration each for 50 ns. The final MD run was carried out for 100 ns (with two replicates for each system with different initial velocities) using charmm36m force-field. All the downstream analysis was carried out using different modules of Gromacs, PYMOL and VMD.

#### Protein-Membrane Interaction

The *g_mindist* command was used to calculate the minimum distance between the protein and the bilayer throughout the simulations.

#### Interacting residues

The distance of residues that interact with membrane was calculated using the n_index and g_minddist module of Gromacs.

#### Video-files for Molecular dynamics simulation

The video files were generated using the PYMOL tool (https://pymol.org/2/).

### Normal mode analysis to understand inter-domain movements

Normal modes to understand inter-domain movements in the RDGB protein were calculated using the ANM 2.1 server. ANM (Anisotropic network model) is a simple NMA tool for analysis of vibrational motions in molecular systems (Eyal et al., 2015). Elastic Network methodology was used and it helped to represent the system at the residue level. The macromolecule is represented as a network of atoms. In the model each protein node is the C_α_ atom of a residue and the overall potential is simply the sum of harmonic potentials between interacting nodes. The network included all interactions within a cut-off distance (distance cut-off of 15 Å). This was the predetermined parameter in the model. Information about the orientation of each interaction with respect to the global coordinates system was considered along the force constant. The force constant was described by Hessian matrix. Each element of the matrix is interaction between two nodes i and j (two C-alpha atoms of two amino-acids). The distance between two nodes was added as a weight at each element of the matrix. The eigen vectors of the matrix describes the vibrational direction and the relative amplitude in the different modes. The mean square fluctuations of individual residues were obtained by summing the fluctuations in the individual modes. 20 normal modes were calculated using the method described above for each of the protein structural models - RDGB, RDGB^(USR1-LNS2)Δ^ andRDGB^LNS2Δ^. The theoretical B-factors were calculated for all modes. Informative modes with the lowest frequency possible and modes responsible for the conformational changes were analyzed further for domain movements. The deformation energy was calculated and modes with least deformation energies along with above mentioned criteria were selected.

**Video files (1-3): Molecular dynamics simulations of protein-membrane system.**

The molecular dynamics simulation was performed using Gromacs for RDGB (video file 1), RDGB^KK/AA^ (video file 2) and RDGB^LNS2Δ^ (video file 3) protein in the presence of DPPC membrane. The simulations were carried out for 100 ns (in replicates) and the movie file was generated using the ‘trjconv’ command of Gromacs. The video files show the interaction between protein and membrane during the entire course of simulation. The protein domains are colour coded as in Figure 1C while the membrane is represented as spheres in grey colour. The residues K1186/A1186 and K1187/A1187 are represented in cyan colour (sphere representation). YW motif is coloured black. The files were generated using PYMOL and can be viewed using a video player with the repeat/loop mode ON.

**Video file 1: Molecular dynamics of RDGB and DPPC membrane**

The residues K1186 and K1187 represented in cyan spheres are seen to interact with the membrane for 80% of the simulation.

**Video file 2: Molecular dynamics of RDGB^KK/AA^ and DPPC membrane**

The residues A1186 and A1187 represented in cyan do not interact with membrane after the system stabilizes and move to a distance greater than 10 Å as the simulation progresses and stabilizes.

**Video file 3: Molecular dynamics of RDGB^LNS2Δ^ and DPPC membrane**

The protein does not form any stable interaction with the membrane during the course of simulation indicating that the LNS2 domain is required for membrane anchoring of the protein.

**Video files (4-6): Normal model analysis of RDGB**

Normal mode analysis (NMA) was carried for RDGB, RDGB^LNS2Δ^ and RDGB^(USR1-LNS2)Δ^ protein to understand inter-domain movements leading to domain interaction. PITPd is marked in red, FFAT motif in green, USR1 in yellow, DDHD domain in blue and LNS2 domain in orange. The ‘YW’ amino acid pair important for membrane docking is marked in black. The videos were generated using PYMOL (educational version) at a speed of 5 FS and with complete smoothening of the trajectory.

**Video file 4: Normal model analysis of RDGB protein:**

The PITPd, FFAT motif and YW motif remain static throughout the NMA, while the DDHD and LNS2 domains move towards each other indicating interaction between the domains.

**Video file 5: Normal model analysis of RDGB^LNS2Δ^ protein:**

The DDHD domain moves towards the PITPd in the absence of the LNS2 domain. This movement is larger compared to the movements observed in wild type protein. The DDHD domain movements are restricted due to the presence of LNS2 domain in the wild type. The YW motif also moves inwards due to the movement of PITPd and DDHD domain. These movement results in the functional consequence as observed in the *in vivo* results.

**Video file 6: Normal model analysis of RDGB^(USR1-LNS2) Δ^ protein**

PITPd and FFAT motif do not show any movements (as in wild type RDGB) in the absence of both the C-terminal domains. The YW motif is restricted in its movements.

---

**Supplementary Figure 1: Molecular dynamics simulation reveals binding energy of RDGB constructs with dVAP-A in the following order: RDGB <RDGB^(DDHD-LNS2)Δ^<RDGB^(USR1-LNS2)Δ^**

Molecular dynamics (MD) simulation was carried out using Desmond module of Schrodinger suite. The final plots with the following parameters for 100 ns MD run (averaged over replicates) have been represented– Total Energy(E), Potential Energy (E_p), Pressure (P), Temperature (T) and Volume (V). The energies are in the order: RDGB/dVAP-A< RDGB^(DDHD-LNS2)Δ^/dVAP-A <RDGB^(USR1-LNS2)Δ^/dVAP-A complex. The X-axis represents the simulation time (ps) and Y-axis represents the Total Energy(E), Potential Energy (E_p), Pressure (P), Temperature (T) and Volume (V) values.

**Supplementary Figure 2: LNS2 domain of RDGB binds lipids at the plasma membrane**

A. Western blot of head extracts made from flies of the mentioned genotype. Tubulin is used as a loading control (N=3). The blot is probed with antibody to GFP. Represented along with it is the LNS2::GFP construct with a flexible Gly/Ser linker (shown in cyan) in between. [Domain structure of LNS2::GFP drawn using Illustrator for Biological Sequences (IBS) software; http://ibs.biocuckoo.org/].

B. Confocal images of S2R+ cells transfected with pJFRC-GFP or pJFRC-LNS2::GFP. UTC: untransfected cells. Green represents GFP. Scale bar= 5 µm.

C. PIP-Strip membranes were incubated over night with S2R+ cell lysate expressing LNS2::GFP and GFP for control. Binding is detected using GFP antibody [LPA= Lysophosphatidic acid, LPC= Lysophosphatidylcholine, PI= Phosphatidylinositol, PI3P= Phosphatidylinositol 3-phosphate, PI4P= Phosphatidylinositol 4-phosphate, PI5P= Phosphatidylinositol 5-phosphate, PE= Phosphatidylethanolamine, PC= Phosphatidylcholine, PS=Phosphatidylserine, PA= Phosphatidic acid, PI(3,4,5)P_3_= Phosphatidylinositol (3,4,5)-trisphosphate, PI(4,5)P_2_= Phosphatidylinositol (4,5)-bisphosphate, PI(3,5)P_2_= Phosphatidylinositol (3,5)-bisphosphate, PI(3,4)P_2_= Phosphatidylinositol (3,4)-bisphosphate, S1P= Sphingosine-1-phosphate].

**Supplementary Figure 3: Charged residues in the LNS2 domain impart its membrane binding property**

A. The minimum distance between RDGB protein (system 1) and the DPPC membrane during the 100 ns molecular dynamics simulation (averaged over replicates) is represented. The X-axis represents simulation time (ps) and Y-axis represents distance between atoms (nm). The minimum distance between the protein and the membrane is 0.2 nm (2 Å) and remains stable throughout the simulation. The graphs were generated using xmgrace module of Gromacs.

B-C. The minimum distance between the residues K1186 and K1187 of RDGB and the DPPC molecule of the membrane is represented (averaged over 100 ns) in figures B and C respectively. The X-axis represents simulation time (ps) and Y-axis represents distance between atoms (nm). Both the residues interact for more than 80% of the simulation with DPPC molecule at a distance of 2 Å. The graphs were generated using xmgrace module of Gromacs.

D. The minimum distance between RDGB^KK/AA^ (system 2) and the DPPC membrane during the 100 ns molecular dynamics simulation (averaged over replicates) is represented. The minimum distance between the protein and the membrane is 0.2 nm (2 Å) at the start of the simulation but increases after 50% of the simulation indicating loss of stable interaction between protein and membrane. The graphs were generated using xmgrace module of Gromacs. The X-axis represents simulation time (ps) and Y-axis represents distance between atoms (nm).

E-F. The minimum distance between the residues A1186 and A1187 of RDGB^KK/AA^ and the DPPC molecule of the membrane is represented in figures E and F respectively (averaged over 100 ns). Both the residues interact at a distance of 3 Å for 50% of the simulation with DPPC molecule at a distance and then move upto 10 Å away from the DPPC molecule indicating loss of interaction between the residues and DPPC molecule. The graphs were generated using xmgrace module of Gromacs. The X-axis represents simulation time (ps) and Y-axis represents distance between atoms (nm).

G. The minimum distance between RDGB^LNS2Δ^ (system 3) and the DPPC membrane during the 100 ns molecular dynamics simulation (averaged over replicates) is represented. The minimum distance between the protein and the membrane is 0.8 nm (8 Å) at the start of the simulation and increases further during the simulation indicating no stable interaction between protein and membrane. The graphs were generated using xmgrace module of Gromacs. The X-axis represents simulation time (ps) and Y-axis represents distance between atoms (nm).

H. Confocal images of S2R+ cells transfected with pJFRC-GFP, pJFRC-LNS2::GFP or pJFRC-LNS2^KK/AA^::GFP. UTC: untransfected cells. Green represents GFP. Scale bar= 5 µm.

## Graphical Abstract

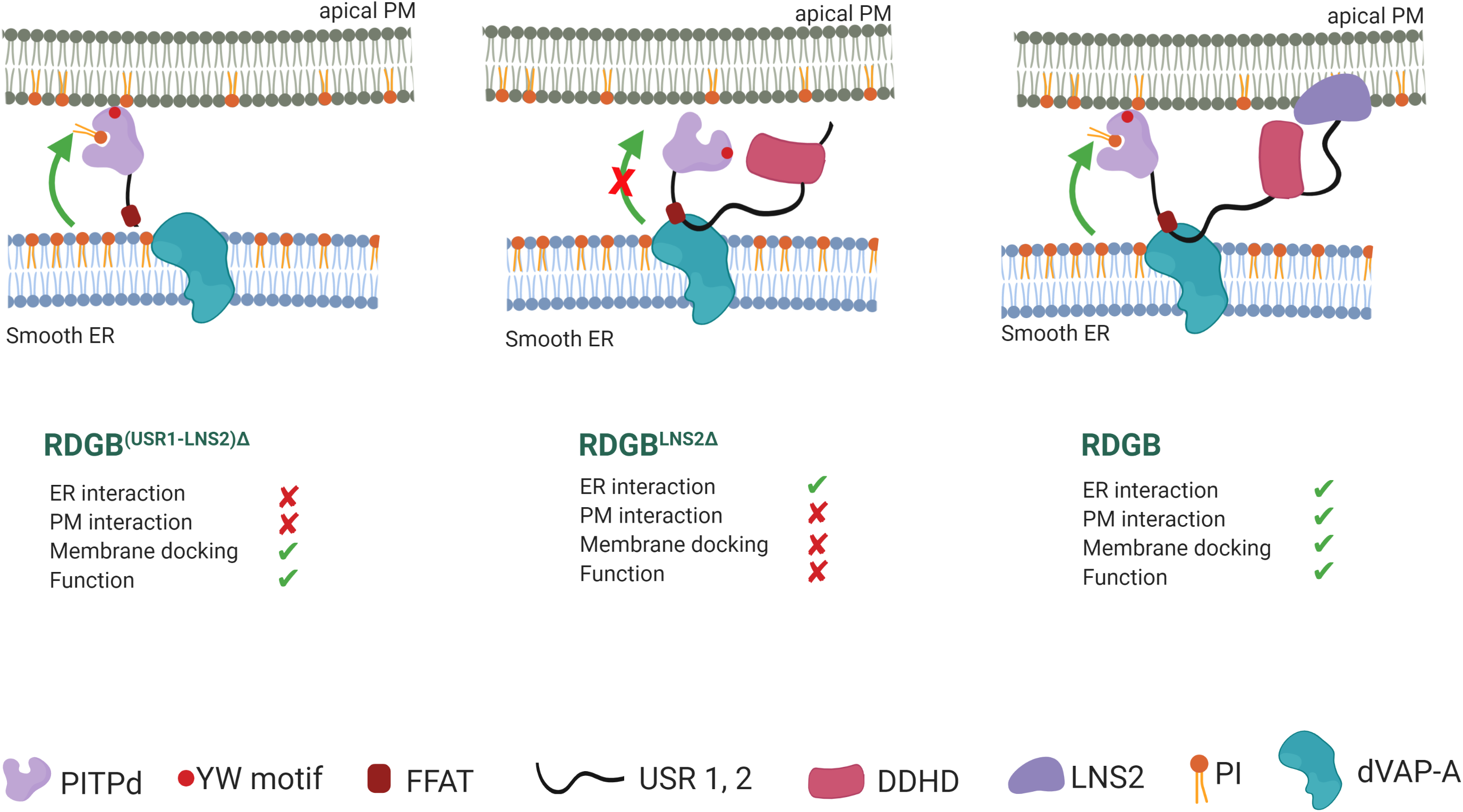

## References

Alli-Balogun, G.O., and Levine, T.P. (2019). Regulation of targeting determinants in interorganelle communication. Curr. Opin. Cell Biol. 57, 106–114.

Ambrish Roy, Alper Kucukural, and & Yang Zhang (2010). I-TASSER: a unified platform for automated protein structure and function prediction. Nat. Protoc. 5, 725–738.

Berendsen, H.J.C., van der Spoel, D., and van Drunen, R. (1995). GROMACS: A message-passing parallel molecular dynamics implementation. Comput. Phys. Commun. 91, 43–56.

Bowers, K.J., Sacerdoti, F.D., Salmon, J.K., Shan, Y., Shaw, D.E., Chow, E., Xu, H., Dror, R.O., Eastwood, M.P., Gregersen, B.A., et al. (2006). Molecular dynamics---Scalable algorithms for molecular dynamics simulations on commodity clusters. In Proceedings of the 2006 ACM/IEEE Conference on Supercomputing-SC ’06, p.

Chakrabarti, P., Kolay, S., Yadav, S., Kumari, K., Nair, A., Trivedi, D., and Raghu, P. (2015). A dPIP5K dependent pool of phosphatidylinositol 4,5 bisphosphate (PIP2) is required for G-protein coupled signal transduction in Drosophila photoreceptors. PLoS Genet. 11, e1004948.

Chen, Y.-J., Quintanilla, C.G., and Liou, J. (2019). Recent insights into mammalian ER–PM junctions. Curr. Opin. Cell Biol. 57, 99–105.

Cockcroft, S., and Raghu, P. (2018). Phospholipid transport protein function at organelle contact sites. Curr. Opin. Cell Biol. 53, 52–60.

Cockcroft, S., Garner, K., Yadav, S., Gomez-Espinoza, E., and Raghu, P. (2016). RdgBα reciprocally transfers PA and PI at ER-PM contact sites to maintain PI(4,5)P2 homoeostasis during phospholipase C signalling in Drosophila photoreceptors. Biochem. Soc. Trans. 44, 286–292.

Cohen, S., Valm, A.M., and Lippincott-Schwartz, J. (2018). Interacting organelles. Curr. Opin. Cell Biol. 53, 84–91.

Dickeson, S.K., Lim, C.N., Schuyler, G.T., Dalton, T.P., Helmkamp Jr., G.M., and Yarbrough, L.R. (1989). Isolation and sequence of cDNA clones encoding rat phosphatidylinositol transfer protein. 264, 16557–16564.

Eyal, E., Lum, G., and Bahar, I. (2015). The anisotropic network model web server at 2015 (ANM 2.0). Bioinformatics 31, 1487–1489.

Fiser, A., and Šali, A. (2003). Modeller: Generation and Refinement of Homology-Based Protein Structure Models. In Methods in Enzymology, pp. 461–491.

Gatta, A.T., and Levine, T.P. (2017). Piecing Together the Patchwork of Contact Sites. Trends Cell Biol. 27, 214–229.

Harris, W.A., and Stark, W.S. (1977). Hereditary retinal degeneration in Drosophila melanogaster. A mutant defect associated with the phototransduction process. J.Gen.Physiol 69, 261–91.

Jamecna, D., Polidori, J., Mesmin, B., Dezi, M., Levy, D., Bigay, J., and Antonny, B. (2019). An Intrinsically Disordered Region in OSBP Acts as an Entropic Barrier to Control Protein Dynamics and Orientation at Membrane Contact Sites. Dev. Cell 49, 220–234.e8.

John, B., and Sali, A. (2003). Comparative protein structure modeling by iterative alignment, model building and model assessment. Nucleic Acids Res. 31, 3982–3992.

Kaiser, S.E., Brickner, J.H., Reilein, A.R., Fenn, T.D., Walter, P., and Brunger, A.T. (2005). Structural basis of FFAT motif-mediated ER targeting. Structure 13, 1035–1045.

Kelley, L.A., Mezulis, S., Yates, C.M., Wass, M.N., and Sternberg, M.J.E. (2015). The Phyre2 web portal for protein modeling, prediction and analysis. Nat. Protoc. 10, 845–858.

Laskowski, R.A., MacArthur, M.W., Moss, D.S., and Thornton, J.M. (1993). PROCHECK: a program to check the stereochemical quality of protein structures. J. Appl. Crystallogr. 26, 283–291.

Milligan, S.C., Alb, J.G., Elagina, R.B., Bankaitis, V.A., and Hyde, D.R. (1997). The phosphatidylinositol transfer protein domain of Drosophila retinal degeneration B protein is essential for photoreceptor cell survival and recovery from light stimulation. J. Cell Biol. 139, 351–363.

Murphy, S.E., and Levine, T.P. (2016). VAP, a Versatile Access Point for the Endoplasmic Reticulum: Review and analysis of FFAT-like motifs in the VAPome. Biochim. Biophys. Acta 1861, 952–961.

Raghu, P., Yadav, S., Babu, N., and Mallampati, N. (2012). Biochimica et Biophysica Acta Lipid signaling in Drosophila photoreceptors. BBA - Mol. Cell Biol. Lipids 1821, 1154–1165.

Saheki, Y., and Camilli, P. De (2017). Endoplasmic Reticulum – Plasma Membrane Contact Sites. Annu. Rev. Biochem. 86, 659–684.

Sanxaridis, P.D., Cronin, M.A., Rawat, S.S., Waro, G., Acharya, U., and Tsunoda, S. (2007). Light-induced recruitment of INAD-signaling complexes to detergent-resistant lipid rafts in Drosophila photoreceptors. Mol. Cell. Neurosci. 36, 36–46.

Schymkowitz J, Borg J, Stricher F, Nys R, Rousseau F, Serrano L. (2005). The FoldX web server: an online force field. Nucleic Acids Res. 33(Web Server issue), W382–W388.

Shadan, S., Holic, R., Carvou, N., Ee, P., Li, M., Murray-Rust, J., and Cockcroft, S. (2008). Dynamics of lipid transfer by phosphatidylinositol transfer proteins in cells. Traffic 9, 1743–1756.

Sukhwal, A., and Sowdhamini, R. (2013). Oligomerisation status and evolutionary conservation of interfaces of protein structural domain superfamilies. Mol. Biosyst. 9, 1652–1661.

Tovchigrechko, A., and Vakser, I.A. (2005). Development and testing of an automated approach to protein docking. Proteins Struct. Funct. Genet. 60, 296–301.

Vihtelic, T.S., Goebl, M., Milligan, S., O’Tousa, J.E., and Hyde, D.R. (1993). Localization of Drosophila retinal degeneration B, a membrane-associated phosphatidylinositol transfer protein. J. Cell Biol. 122, 1013–1022.

Yadav, S., Garner, K., Georgiev, P., Li, M., Gomez-Espinosa, E., Panda, A., Mathre, S., Okkenhaug, H., Cockcroft, S., and Raghu, P. (2015). RDGBα, a PtdIns-PtdOH transfer protein, regulates G-protein coupled PtdIns(4,5)P2 signalling during Drosophila phototransduction. J. Cell Sci. 128, 3330–3344.

Yadav, S., Cockcroft, S., and Raghu, P. (2016). The Drosophila photoreceptor as a model system for studying signalling at membrane contact sites. Biochem. Soc. Trans. 44, 447–451.

Yadav, S., Thakur, R., Georgiev, P., Deivasigamani, S., Krishnan, H., Ratnaparkhi, G., and Raghu, P. (2018). RDGBα localization and function at membrane contact sites is regulated by FFAT–VAP interactions. J. Cell Sci. 131, 1.

