## Supplemental Data for "The lipid transfer function of RDGB at ER-PM contact sites is regulated by multiple interdomain interactions"

**RDGB/dVAP-A**

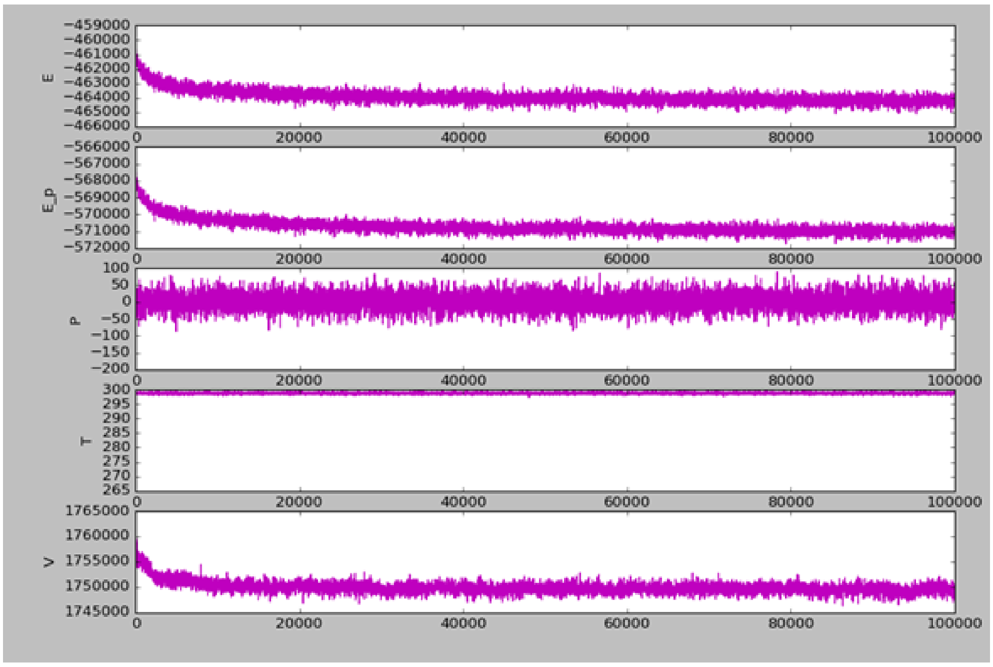


**RDGB$^{(DDHD\text{-}LNS2)\Delta}$/dVAP-A**

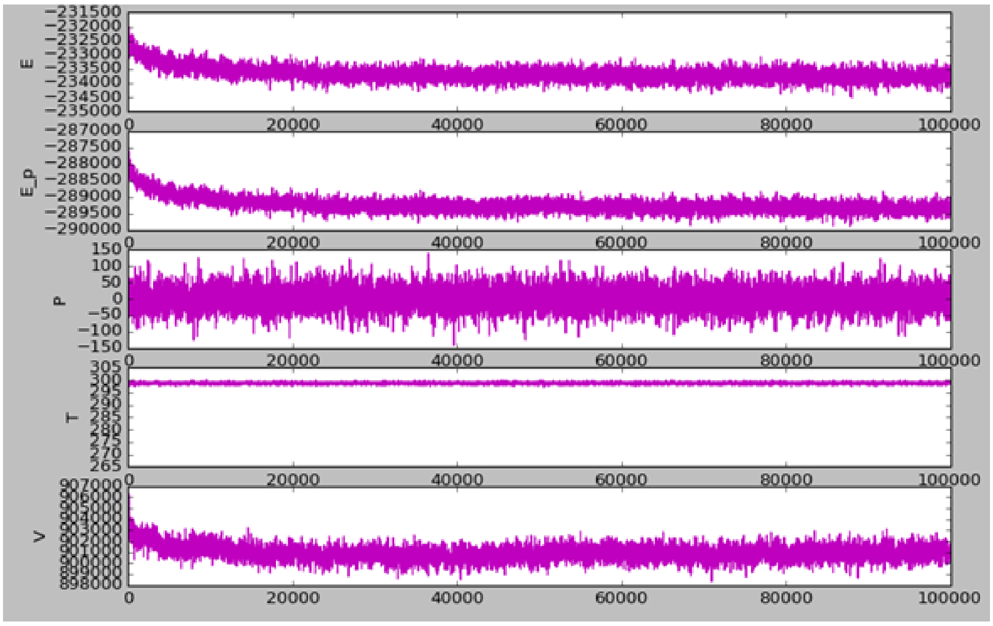


**RDGB$^{(USR1\text{-}LNS2)\Delta}$/dVAP-A**

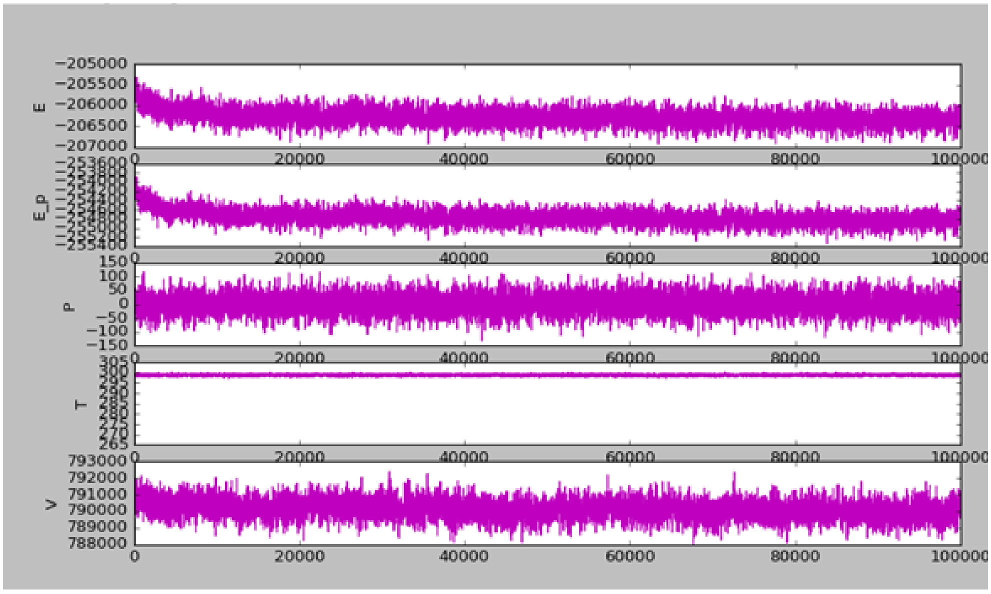

A.

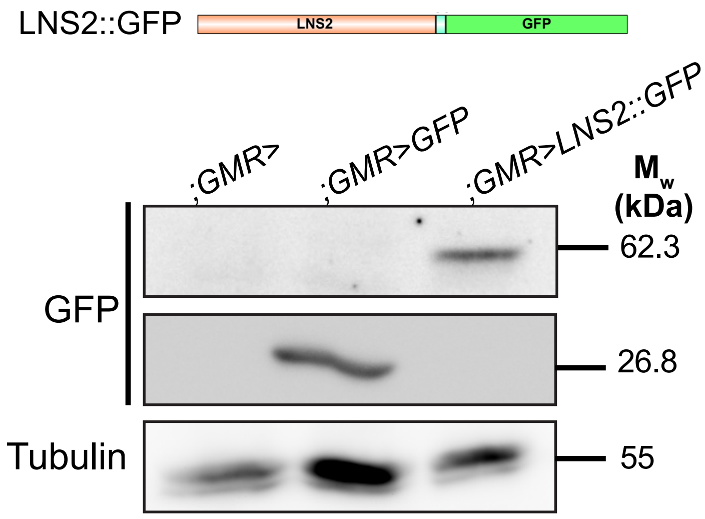
LNS2::GFP
LNS2
GFP
;GMR>
;GMR>GFP
;GMR>LNS2::GFP
$M_w$ (kDa)
GFP
62.3
26.8
Tubulin
55

B.

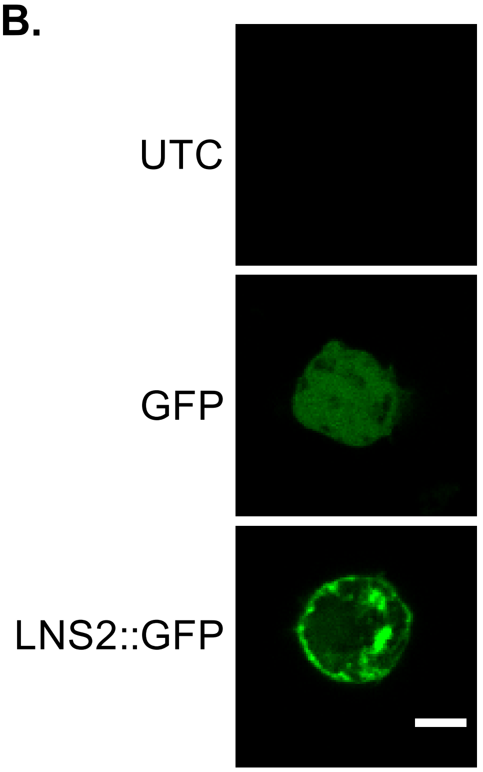
UTC
GFP
LNS2::GFP

C.

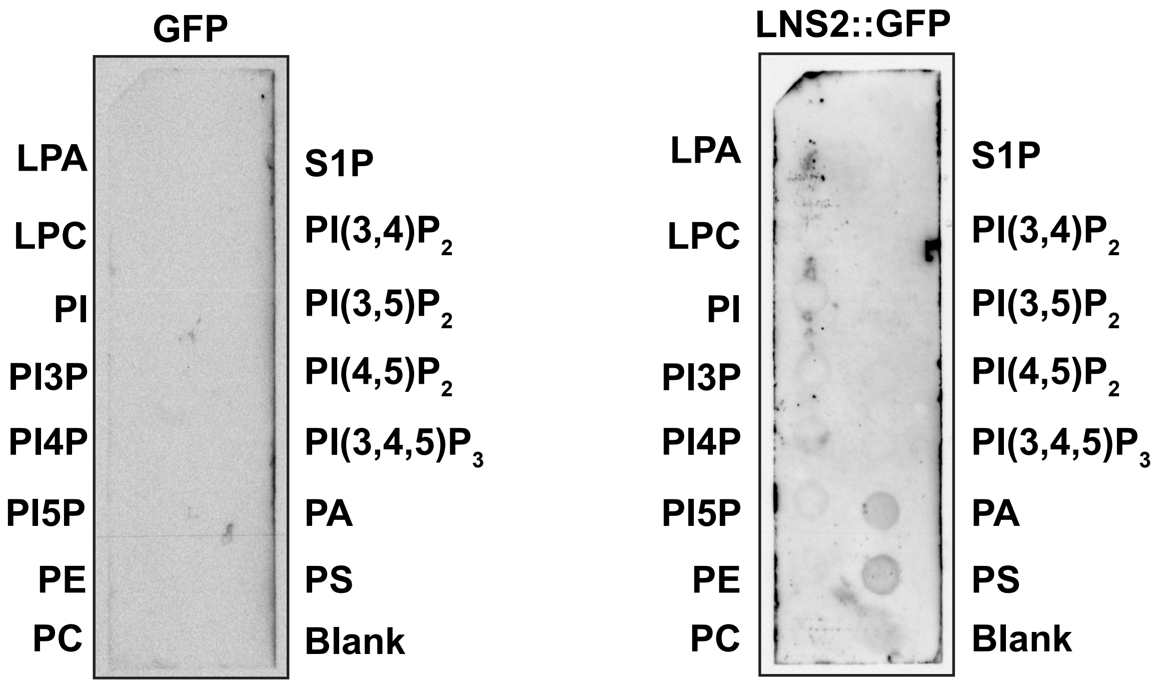
GFP
LNS2::GFP
LPA
LPC
PI
PI3P
PI4P
PI5P
PE
PC
S1P
$PI(3,4)P_2$
$PI(3,5)P_2$
$PI(4,5)P_2$
$PI(3,4,5)P_3$
PA
PS
Blank

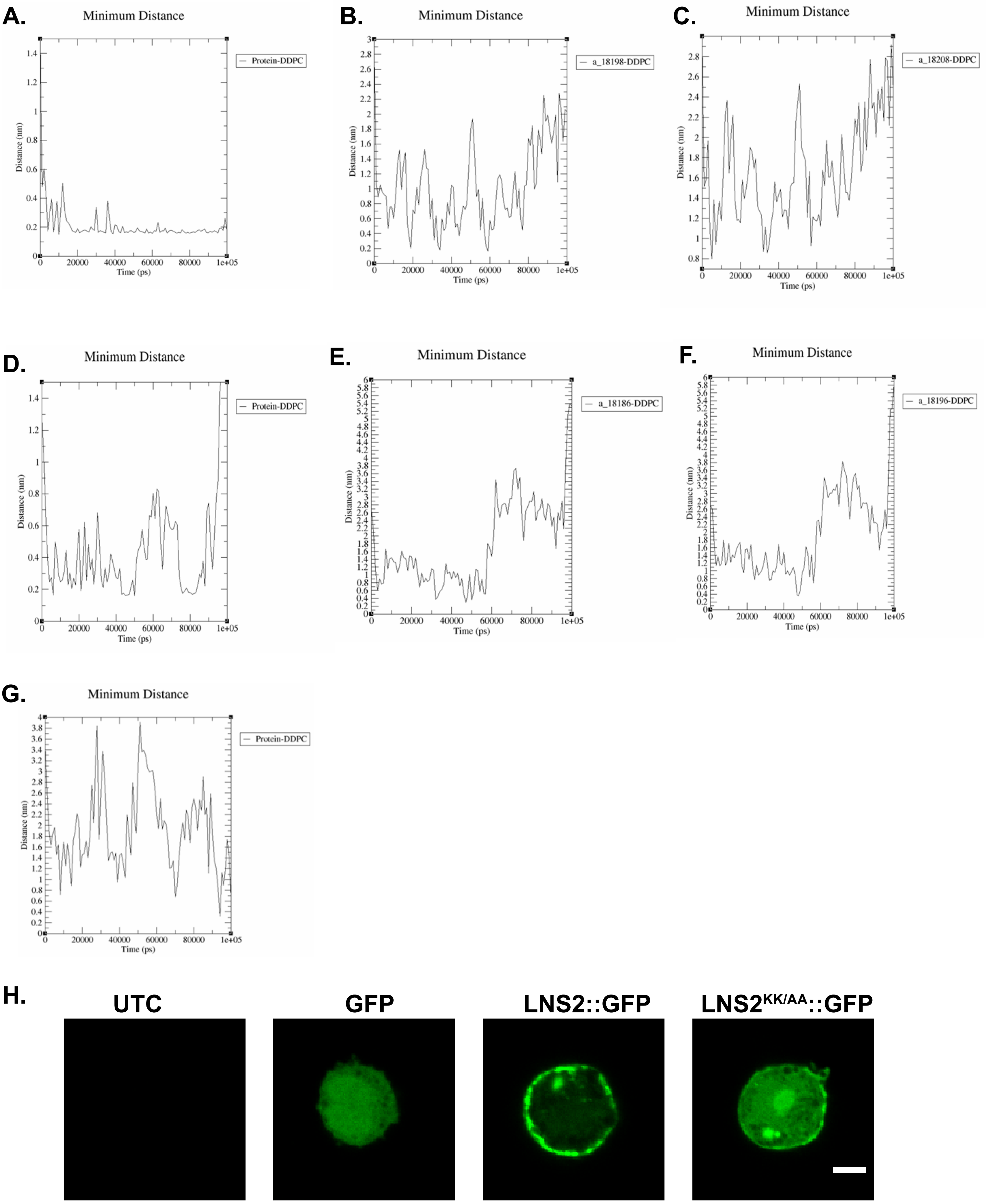


Basak et al., Supplemental Data 3
